# Ipsilateral or contralateral boosting of mice with mRNA vaccines confers equivalent immunity and protection against a SARS-CoV-2 Omicron strain

**DOI:** 10.1101/2024.04.05.588051

**Authors:** Baoling Ying, Chieh-Yu Liang, Pritesh Desai, Suzanne M. Scheaffer, Sayda M. Elbashir, Darin K. Edwards, Larissa B. Thackray, Michael S. Diamond

**Affiliations:** Department of Medicine, Washington University School of Medicine, St. Louis, MO 63110, USA; Department of Pathology & Immunology, Washington University School of Medicine, St. Louis, MO, USA; Moderna, Inc., Cambridge MA, USA; Department of Molecular Microbiology, Washington University School of Medicine, St. Louis, MO, USA; The Andrew M. and Jane M. Bursky Center for Human Immunology and Immunotherapy Programs, Washington University School of Medicine. St. Louis, MO, USA; Center for Vaccines and Immunity to Microbial Pathogens, Washington University School of Medicine, Saint Louis, MO, USA

## Abstract

Boosting with mRNA vaccines encoding variant-matched spike proteins has been implemented to mitigate their reduced efficacy against emerging SARS-CoV-2 variants. Nonetheless, in humans, it remains unclear whether boosting in the ipsilateral or contralateral arm with respect to the priming doses impacts immunity and protection. Here, we boosted K18-hACE2 mice with either monovalent mRNA-1273 (Wuhan-1 spike) or bivalent mRNA-1273.214 (Wuhan-1 + BA.1 spike) vaccine in the ipsilateral or contralateral leg relative to a two-dose priming series with mRNA-1273. Boosting in the ipsilateral or contralateral leg elicited equivalent levels of serum IgG and neutralizing antibody responses against Wuhan-1 and BA.1. While contralateral boosting with mRNA vaccines resulted in expansion of spike-specific B and T cells beyond the ipsilateral draining lymph node (DLN) to the contralateral DLN, administration of a third mRNA vaccine dose at either site resulted in similar levels of antigen-specific germinal center B cells, plasmablasts/plasma cells, T follicular helper cells and CD8^+^ T cells in the DLNs and the spleen. Furthermore, ipsilateral and contralateral boosting with mRNA-1273 or mRNA-1273.214 vaccines conferred similar homologous or heterologous immune protection against SARS-CoV-2 BA.1 virus challenge with equivalent reductions in viral RNA and infectious virus in the nasal turbinates and lungs. Collectively, our data show limited differences in B and T cell immune responses after ipsilateral and contralateral site boosting by mRNA vaccines that do not substantively impact protection against an Omicron strain.

**IMPORTANCE:** Sequential boosting with mRNA vaccines has been effective strategy to overcome waning immunity and neutralization escape by emerging SARS-CoV-2 variants. However, it remains unclear how the site of boosting relative to the primary vaccination series shapes optimal immune responses or breadth of protection against variants. In K18-hACE2 transgenic mice, we observed that boosting with historical monovalent or variant-matched bivalent vaccines in the ipsilateral or contralateral limb elicited comparable levels of serum spike specific antibody and antigen-specific B and T cells responses. Moreover, boosting on either side conferred equivalent protection against a SARS-CoV-2 Omicron challenge strain. Our data in mice suggest that the site of boosting with an mRNA vaccine does not substantially impact immunity or protection against SARS-CoV-2 infection.

## INTRODUCTION

The coronavirus disease (COVID-19) pandemic has caused an estimated 775 million infections and 7.2 million deaths worldwide (https://covid19.who.int/). Multiple vaccines (*e.g*., mRNA, viral-vectored, inactivated, nanoparticle, and protein subunit-based) targeting the SARS-CoV-2 spike protein were deployed with billions of doses now administered globally. Many of these vaccines are effective against the severe disease, hospitalization, and death caused by SARS-CoV-2, with an estimated 20 million lives saved during the first year of vaccine rollout ^1^. Despite the success of the COVID-19 vaccines, multiple waves of SARS-CoV-2 variants (*e.g*., B.1.1.7 [Alpha], B.1.351 [Beta], B.1.617.2 [Delta], BA.1, BA.5, XBB.1.5 and currently circulating JN.1 [Omicron]) continue to emerge, each with different constellations of amino acid substitutions, deletions, and insertions in the spike protein and elsewhere in the genome. As these successive variants often show increased transmissibility and greater antibody escape potential, they pose challenges to the efficacy of existing vaccines. Indeed, the antigen-shifted Omicron variants compromised the protective efficacy of immune responses elicited by vaccination or prior infection with non-Omicron strains ^2–5^.

Beyond changes in the spike protein of variants, waning immunity poses a separate challenge to vaccine efficacy. Although mRNA vaccines (Moderna mRNA-1273; Pfizer-BioNTech BNT162b2) induce durable germinal center B cells (GCB) responses for at least 6 months ^6–9^, the serum neutralizing antibody responses wane within 3 to 6 months and decline further by 8 months, with an estimated half-life of approximately 60 days ^10–12^. Waning humoral antibody and emergence of resistant, highly transmissible variants correlate with an increased frequency of breakthrough infections in vaccinated individuals ^13,14^.

To address concerns arising from reduced efficacy of vaccines against current and future variants, third, fourth, fifth, and even sixth doses (herein, boosters) of monovalent (Wuhan-1 or more recently, XBB.1.5) or bivalent mRNA vaccines encoding the historical (Wuhan-1) and variant-matched (BA.1 or BA.5) spike protein have been implemented. Indeed, mRNA booster vaccines have been shown to increase the magnitude and breadth of neutralizing antibodies against SARS-CoV-2 ^15–20^. One frequent question regarding boosting is which arm should be used and whether the site injected affects immunogenicity and protection with mRNA or other COVID-19 vaccines. A study in mice with alum-adjuvanted influenza protein antigens showed that ipsilateral boosting with a homologous vaccine induced recall germinal center responses with B cells having greater levels of somatic hypermutation and antibodies with higher avidity and cross-reactivity than boosting at a remote site ^21^. In comparison, data from human studies with Pfizer-BioNTech BNT162b2 mRNA vaccine are less certain, as two studies reached different conclusions. Whereas Ziegler *et al* reported that ipsilateral boosting with homologous BNT162b2 mRNA vaccine elicited higher serum neutralizing titers two weeks later compared to contralateral boosting ^22^, Fazli *et al* showed that contralateral boosting with a homologous mRNA vaccine was associated with higher binding and neutralizing antibody titers at 8- and 14-months ^23^. Moreover, a study in mice with a two-dose regimen of BNT162b2 mRNA vaccine reported that ipsilateral compared to contralateral leg boosting produced higher numbers of GCBs that bound the receptor binding domain (RBD) of an ancestral strain in the draining lymph node (DLN) and higher levels of antibody with enhanced affinity against RBD of ancestral and Omicron strains but only at the early (day 9) but not late (week 19) time points ^24^. However, the impact of ipsilateral versus contralateral boosting on T cell responses or protective immunity in the context of homologous or heterologous SARS-CoV-2 challenge has not been tested.

Here, we evaluated the impact of ipsilateral and contralateral leg boosting with preclinical versions of historical (mRNA-1273, Wuhan-1 spike) and bivalent (mRNA-1273.214, Wuhan-1 + BA.1 spike) mRNA vaccines on adaptive B and T cell responses and protection against challenge with a BA.1 variant in susceptible K18-hACE2 transgenic mice. We observed that ipsilateral and contralateral boosting elicited comparable levels of serum spike specific antibody as well as antigen-specific B and T cells responses. The differential impact of boosting site on protective response in the context of virus challenge was limited, as ipsilateral and contralateral boosting with either mRNA-1273 or mRNA-1273.214 vaccines conferred similar levels of virological protection against BA.1 infection.

## RESULTS

### Antibody responses after boosting with mRNA vaccines

To compare serum antibody responses following vaccine boosting on the ipsilateral or contralateral side, cohorts of 7- to 9-week-old female K18-hACE2 transgenic mice were immunized intramuscularly in the left hind leg twice over a 3 week-interval with a preclinical version of mRNA-1273 (**Fig 1A**) that encodes for the prefusion-stabilized spike protein of SARS-CoV-2 Wuhan-1 strain ^25^. Animals were boosted 12 weeks later in the corresponding site in the left (ipsilateral) or right (contralateral) leg with 0.25 µg of mRNA-1273 or a preclinical bivalent mRNA-1273.214 vaccine comprised of 1:1 mixture of mRNA-1273 and the BA.1-matched mRNA-1273.529 vaccine (**Fig 1A**). Boosting with these two different vaccines enabled analysis of the effects of immunization site on the breadth of the immune response. Serum was collected one day before and 4 weeks after boosting, and IgG responses against recombinant Wuhan-1 and BA.1 pre-fusion-stabilized spike (S-2P) (**Fig 1B-E**) or RBD (**Fig 1F-I**) proteins were determined by ELISA. Prior to boosting, as expected, similar serum IgG titers were detected among all four groups against spike of WA1/2020 D614G spike (**Fig 1B**), and the levels against BA.1 spike were approximately 7- to 8-fold lower (p < 0.01) than against Wuhan-1 spike (**Fig 1C**). Boosting with either mRNA-1273 or mRNA-1273.214 elicited enhanced IgG responses against the spike (**Fig 1B-E**) and RBD (**Fig 1F-I**) of Wuhan-1 and BA.1. For both mRNA-1273 and mRNA-1273.214 boosts, lower serum IgG titers were observed against the BA.1 spike (4 to 5-fold, p < 0.01) and RBD (17 to 20-fold, p < 0.0001) than the corresponding Wuhan-1 proteins. In pre- and post-boost comparisons, boosting with mRNA-1273 increased IgG binding titers against Wuhan-1 spike (8 to 10-fold, p < 0.01, **Fig 1D**) or RBD (6 to-8-fold, p < 0.0001, **Fig 1H**) with smaller increases against BA.1 RBD. In comparison, mRNA-1273.214 induced larger increases against the BA.1 RBD (11 to 15-fold, p < 0.01, **Fig 1I**). In the context of the booster location, ipsilateral and contralateral injections with mRNA-1273 or mRNA-1273.214 induced similar IgG binding titers against either Wuhan-1 or BA.1 protein.

**Figure 1.**
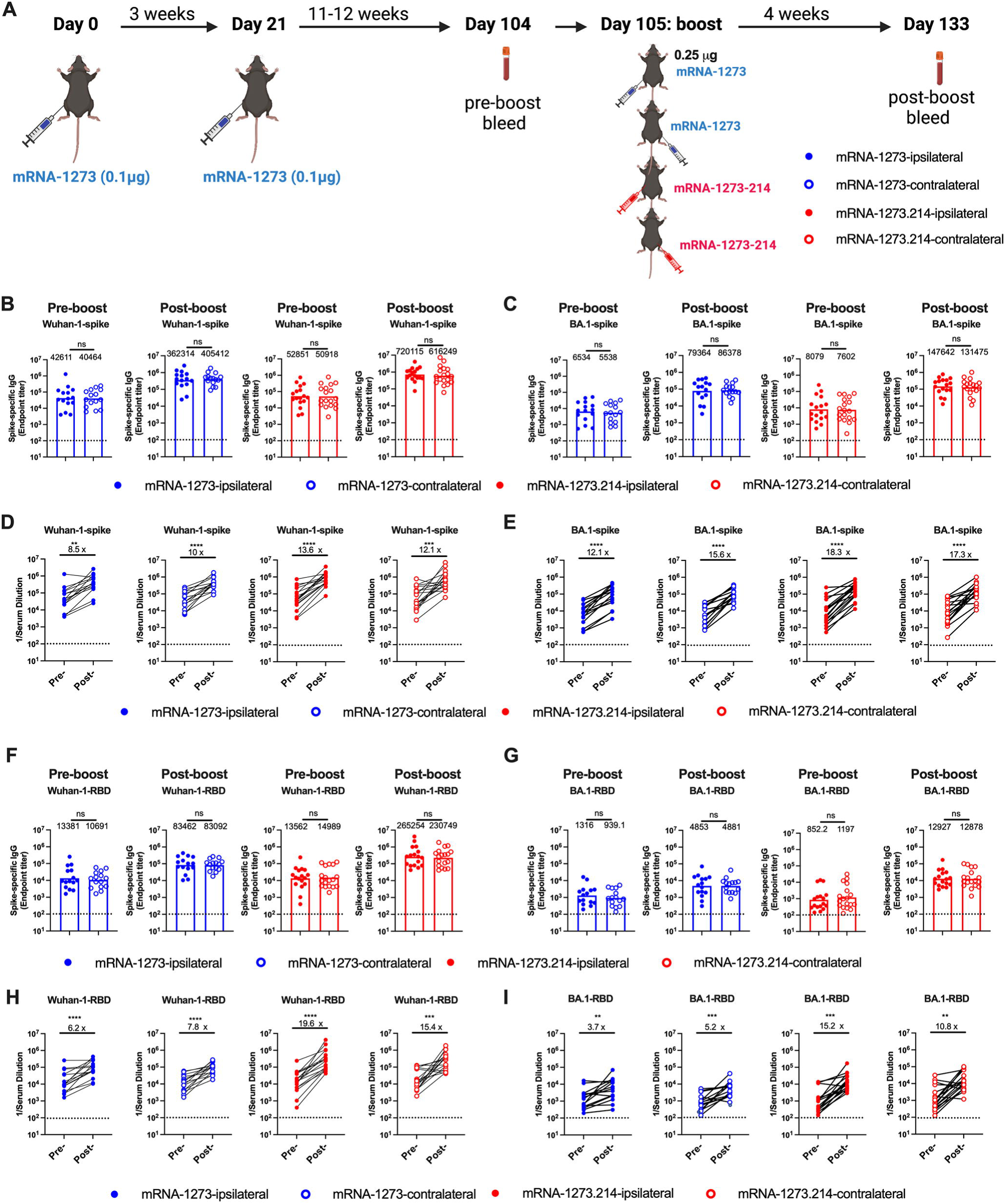
Serum IgG responses in K18-hACE2 mice following boosting. Seven- to nine-week-old female K18-hACE2 mice were immunized with a primary two-dose vaccination series spaced three weeks apart in the left hind leg with 0.1 μg of mRNA-1273. Animals were boosted 11 to 12 weeks later in the left (ipsilateral) or right (contralateral) leg with 0.25 μg of mRNA-1273 or bivalent mRNA-1273.214. One day before (pre-boost) or four weeks after boost (post-boost), serum IgG titers were determined by ELISA. **A**. Scheme of immunizations and blood collection. **B-C**. Serum IgG responses against spike of Wuhan-1(**B**) or BA.1 (**C**). **D-E**. Paired analysis of serum IgG titers of pre- and post- boost against WA1/2020 D614G (**D**) and BA.1 (**E**). **F-G**. Serum IgG responses against RBD domain of Wuhan-1 (**F**) or BA.1 (**G**). **H-I**. Paired analysis of serum IgG titers of pre- and post- boost against WA1/2020 D614G (**H**) and BA.1 (**I**) (n = 15-18, two experiments, column heights indicate geometric mean titers (GMT), dotted lines show the limit of detection [LOD]). GMTs or fold-changes are indicated above corresponding graphs. Statistical analyses: **B, C, F, G**, Mann-Whitney test; **D, E, H, I,** Wilcoxon matched signed-rank test. * p < 0.05; ** p < 0.01, ∗∗∗ p < 0.001, ∗∗∗∗ p < 0.0001.

We next characterized the inhibitory effects of pre- and post-boost serum on SARS-CoV-2 infectivity using an established focus-reduction neutralization test (FRNT) ^26^ and authentic SARS-CoV-2 WA1/2020 D614G and BA.1 viruses (**Fig 2A-D and S1**). Because of limited amounts of sera, we started dilutions at 1/60, which is just above the estimated threshold level of neutralization associated with protection in humans ^27^. As expected, twelve weeks after the primary immunization series with mRNA-1273 and before boosting, the serum neutralizing titers against WA1/2020 D614G were equivalent among all four groups (**Fig 2A**). However, and as observed in previous studies ^16,17,28^, serum from mice receiving two doses of mRNA-1273 showed poor neutralizing activity against BA.1, with most falling below the limit of detection of the assay (**Fig 2B**). One month after boosting with either mRNA-1273 or mRNA-1273.214, serum neutralizing titers against WA1/2020 D614G rose 5 to 8-fold (p < 0.001) (**Fig 2A and C**) with minimal differences observed after ipsilateral or contralateral leg boosting. In comparison, ipsilateral or contralateral boosting with mRNA-1273 showed marginally increased neutralizing titers against BA.1, which did not achieve statistical significance (**Fig 2B and D**). As expected for a variant-matched vaccine, boosting with mRNA-1273.214 induced 3 to 5-fold higher neutralizing responses against BA.1 (p < 0.001), with ipsilateral boosters showing slightly higher titers than contralateral boosters, although these differences did not attain statistical significance (**Fig 2B and D**). Overall, only small differences in neutralizing activity were observed when comparing the site (ipsilateral versus contralateral) of mRNA vaccine boosting.

**Figure 2.**
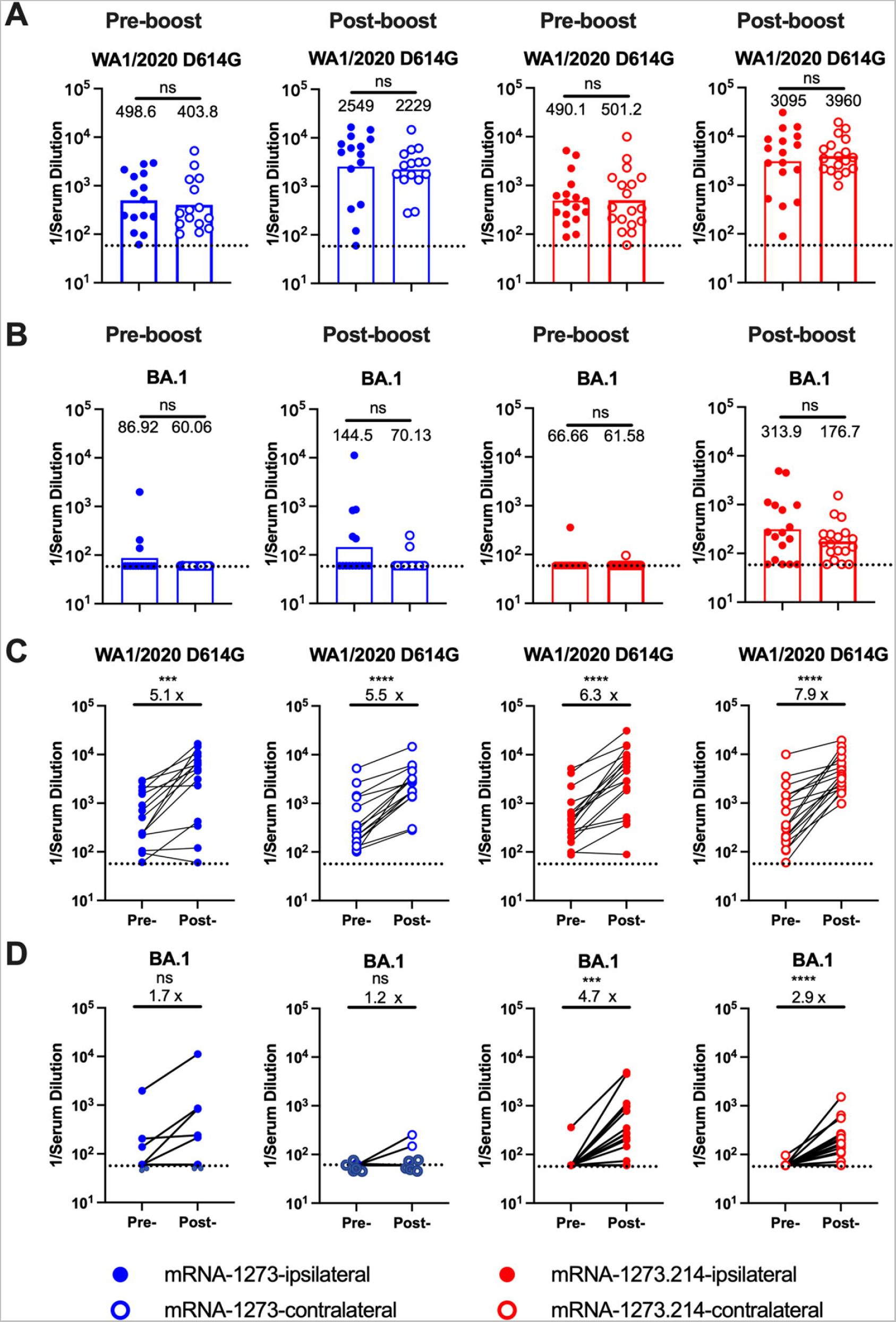
Serum neutralizing antibody responses against WA1/2020 D614G and BA.1. One day before or four weeks after ipsilateral or contralateral hind leg boosting with mRNA-1273 or mRNA-1273.214, serum neutralizing antibody were determined by focus reduction neutralization test (FRNT) against the indicated authentic SARS-CoV-2 strains. **A.** WA1/2020 D614G. **B.** BA.1 (n = 15-18, two experiments, boxes illustrate GMTs, dotted lines show the limit LOD). **C-D**. Paired analysis of serum neutralizing titers of pre- and post- boost against WA1/2020 D614G (**C**) and BA.1 (**D**) (n = 15-18, two experiments, column heights indicate GMTs, dotted lines show the LOD. GMTs or fold-changes are indicated above corresponding graphs). Statistical analyses: **A-B**, Mann-Whitney test; **C-D,** Wilcoxon matched- pairs signed-rank test. ns, not significant; * p < 0.05; ** p < 0.01, ∗∗∗ p < 0.001, ∗∗∗∗ p < 0.0001.

### B cell responses after boosting with mRNA vaccines

To assess further the impact of the site of boosting on immune responses, we interrogated B cells in the DLN and spleen. In this set of experiments, we applied the same vaccination scheme (**Fig 1A**) but boosted with a higher 1 µg dose of mRNA-1273 or bivalent mRNA-1273.214 either on ipsilateral (left) or contralateral (right) side; a higher booster dose was used to enhance our ability to detect spike-specific B cell responses by flow cytometry. Seven days after boosting, we harvested the left and right inguinal DLNs as well as the spleen to characterize GCBs (CD19^+^IgD^lo^GL7^+^Fas^+^) and plasmablast/plasma cell (PB/PC: CD19^+^IgD^lo^TACI^+^CD138^+^) responses (**Fig 3**, **4 and S2**). Ipsilateral and contralateral leg boosting with mRNA-1273 or mRNA-1273.214 vaccines induced 7 to 8-fold higher frequencies (p <0.0001) and 100 to 200-fold higher numbers (p < 0.0001) of GCBs in their respective DLN (ipsilateral, left; contralateral, right) than in naïve mice (**Fig 3A and S3A**). Boosting with either vaccine induced lower GCB responses in the non-draining, contralateral LN with only 2 to 5-fold higher total numbers of GCB compared to unvaccinated mice (**Fig 3A**). These data suggest that mRNA vaccine boosting preferentially induces GC reactions in the DLN, which is consistent with data from human studies ^7,9^. Nonetheless, mRNA-1273 and mRNA-1273.214 vaccines generally elicited similar overall GCB responses after ipsilateral and contralateral boosts.

**Figure 3.**
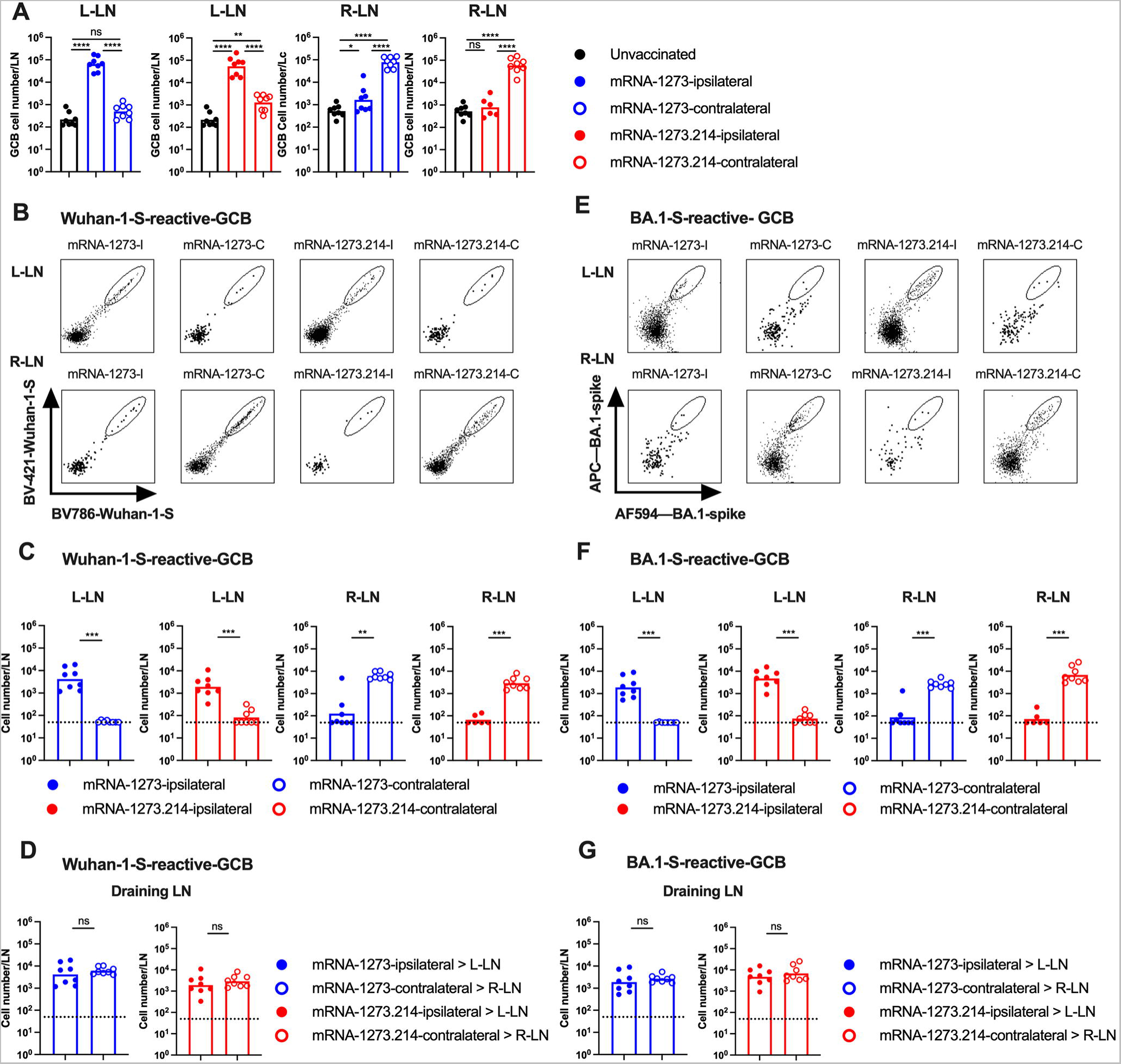
Germinal center B cell responses in the lymph nodes following boosting with mRNA-1273 or mRNA1273.214. Seven- to nine-week-old female K18-hACE2 mice were immunized with a primary two-dose vaccination series spaced three weeks apart in the left hind leg with mRNA-1273. Animals then were boosted 11 to 12 weeks later in the left (ipsilateral) or right (contralateral) leg with 1 µg of mRNA-1273 or mRNA-1273.214. Seven days after boosting, inguinal LNs from the left and right side were analyzed for GCB responses by flow cytometry. **A**. Quantification of total number of CD19^+^IgD^lo^GL7^+^Fas^+^GCBs in respective LNs. **B**. Representative flow cytometry scatter plots of Wuhan-1 spike-reactive GCBs. **C**. Total numbers of Wuhan-1 spike-reactive GCBs in the respective LNs. **D**. Comparison of numbers of Wuhan-1 spike-reactive GCBs in the DLNs. **E**. Representative flow cytometry scatter plots of BA.1 spike- reactive GCBs. **F**. Total number of BA.1 spike-reactive GCBs in respective LNs. **G.** Comparison of numbers of BA.1 spike-reactive GCBs in the DLNs. Data are from two experiments (n = 7-8, each data point represents an individual mouse, column heights indicate geometric mean values, dotted lines show the LOD). Statistical analyses: **A,** One-way ANOVA with Tukey’s post- test; **C, D, F, G,** unpaired two-tailed Mann-Whitney test: ns, not significant; * p < 0.05, ∗∗ p < 0.01, ∗∗∗ p < 0.001, ∗∗∗∗ p < 0.0001.

**Figure 4.**
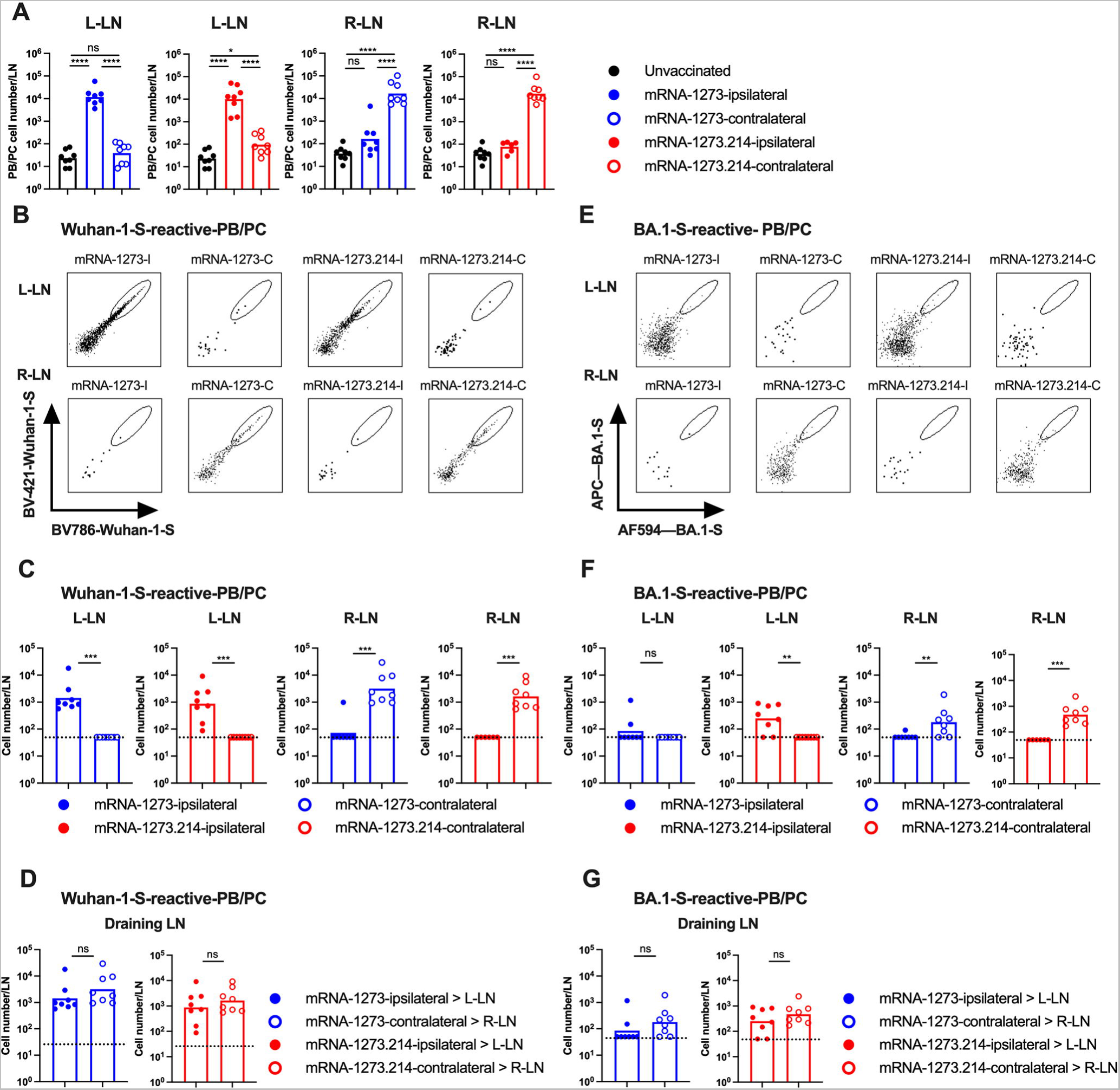
Plasmablast and plasma cell responses in the lymph nodes following boosting with mRNA-1273 or mRNA1273.214. K18-hACE2 mice were immunized and boosted as described in Fig 3. Seven days after boosting, inguinal LNs from the left and right side were analyzed for PB/PC responses by flow cytometry. **A**. Total number CD19^+^IgD^lo^CD138^+^TACI^+^ PBs/PCs. **B**. Representative flow cytometry scatter plots of Wuhan-1 spike-reactive PBs/PCs. **C**. Total number of Wuhan-1 spike-reactive PBs/PCs in respective LNs. **D**. Comparison of numbers of Wuhan-1 spike-reactive PBs/PCs in the DLNs. **E**. Representative flow cytometry scatter plots of BA.1 spike reactive PBs/PCs. **F**. Total number of BA.1 spike reactive PB/PC cells in respective LNs. **G.** Comparison of numbers of BA.1 spike reactive PBs/PCs in the DLNs. Data are from two independent experiments (n = 7-8, each data point represents an individual mouse, column heights indicate geometric mean values, dotted lines show the LOD). Statistical analyses: **A,** One-way ANOVA with Tukey’s post-test; **C, D, F, G,** unpaired two-tailed Mann-Whitney test: ns, not significant; * p < 0.05, ∗∗ p ≤ 0.01, ∗∗∗ p < 0.001, ∗∗∗∗ p < 0.0001.

We next profiled antigen-specific GCB responses using fluorescently labeled Wuhan-1 and BA.1 spike protein and flow cytometry (**Fig 3B-G, and S3B-C**). Ipsilateral and contralateral boosting with mRNA-1273 or mRNA-214 vaccines both elicited robust GCB responses in DLNs against Wuhan-1 (**Fig 3B-D**) and BA.1 (**Fig 3E-G**) spike proteins. Regardless of the boost location or mRNA vaccine used, higher numbers and frequencies of spike-specific GCBs were detected in the DLN on the side of the boost. When we compared the mRNA-1273 and mRNA-1273.214 response in the DLN where the dominant GC reaction occurred, mRNA-1273 elicited approximately 2-fold higher numbers (**Fig 3C**) and frequency (**Fig S3B**) of GCBs that bound Wuhan-1 spike than mRNA-1273.214. In comparison, mRNA-1273.214 elicited approximately 2 to 3-fold higher numbers (**Fig 3F**) and frequency (**Fig S3C**) of GCBs that bound BA.1 spike than mRNA-1273. Ipsilateral and contralateral leg boosting elicited relatively equivalent overall numbers of Wuhan-1-spike reactive or BA.1-spike reactive GCBs in the DLNs regardless of vaccines being used (**Fig 3D and G**), with substantially reduced responses observed in LNs on the side opposite of boosting (**Fig 3C and F**).

We also assessed total and antigen-specific PB/PC responses in the DLNs (**Fig 4 and S3D-F**) and observed a pattern similar to the GCB response. Ipsilateral and contralateral leg boosting with mRNA-1273 or mRNA-1273.214 vaccines elicited greater total (**Fig 4A, S3D**) and spike-specific (**Fig 4B-G and S3E-F**) PB/PC responses in the DLN on the same side of the boost. mRNA-1273 elicited a higher frequency (**Fig S3E**) and number of PB/PCs that bound to Wuhan-1 spike than mRNA-1273.214 (**Fig 4C**), and reciprocally mRNA-1273.214 elicited a greater frequency (**Fig S3F**) and number of PB/PCs that bound to BA.1 spike (**Fig 4F**). In terms of the effect of the boosting site, ipsilateral and contralateral leg boosting elicited relatively similar numbers of Wuhan-1- or BA.1-spike reactive PBs/PCs in the respective DLNs regardless of the vaccine used (**Fig 4D and G**), with lower responses in LNs achieved at the site opposite of boosting (**Fig 4C and F**). Overall, boosting with mRNA vaccines resulted in comparable cellular B cell responses against homologous or heterologous spike antigens in the DLN regardless of whether the boost occurred ipsilateral or contralateral to the site of the primary vaccination series.

We also measured the total and spike-specific PB/PC response in the spleen seven days after boosting (**Fig S4**). Ipsilateral or contralateral boosting with mRNA-1273 or mRNA-1273.214 vaccines induced 5 to 10-fold higher numbers (p <0.0001) of total PB/PC in the spleen than in unvaccinated mice (**Fig S4A**). Mice boosted with either mRNA-1273 vaccines had 6 to 9-fold higher numbers (p < 0.05) of antigen-specific PB/PCs that bound spike of Wuhan-1 compared to BA.1 (**Fig S4B-C**). However, and like we observed in the DLN, ipsilateral or contralateral boosting with mRNA-1273 or mRNA-1273.214 induced equivalent levels of PB/PCs in the spleen that bound Wuhan-1 (**Fig S4B**) and BA.1 (**Fig S4C**) spike proteins.

### T cell responses after boosting with mRNA vaccines

Since T cell immunity also contributes to protection against SARS-CoV-2 infection and disease ^8,29–32^, we interrogated their responses after mRNA vaccine boosting (**Fig 5, S5, and S6**). At 7 days after ipsilateral or contralateral leg boosting with mRNA-1273 or mRNA-1273.214 vaccines, similar or slightly higher numbers of total CD3^+^ T cells were present in the respective DLN (ipsilateral, left; contralateral, right) compared to non-boosted mice (**Fig 5A**). We next evaluated spike-specific CD8^+^ T responses using class I MHC tetramers displaying a conserved immunodominant peptide (S_539-546_; VNFNFNGL). Ipsilateral and contralateral boosting with mRNA-1273 or mRNA-214 vaccines elicited higher frequencies and numbers of tetramer^+^ CD8^+^ T cells in the DLN, with reduced responses observed on the side opposite of boosting (**Fig 5B-D**). Ipsilateral and contralateral mRNA vaccine boosting elicited comparable numbers of tetramer+ CD8^+^ T cells in the DLNs regardless of the vaccine used (**Fig 5E**). Ipsilateral or contralateral boosting with mRNA-1273 or mRNA-1273.214 mRNA also elicited comparable frequencies and numbers of tetramer+ CD8^+^ T cells in the spleen (**Fig 5F**).

**Figure 5.**
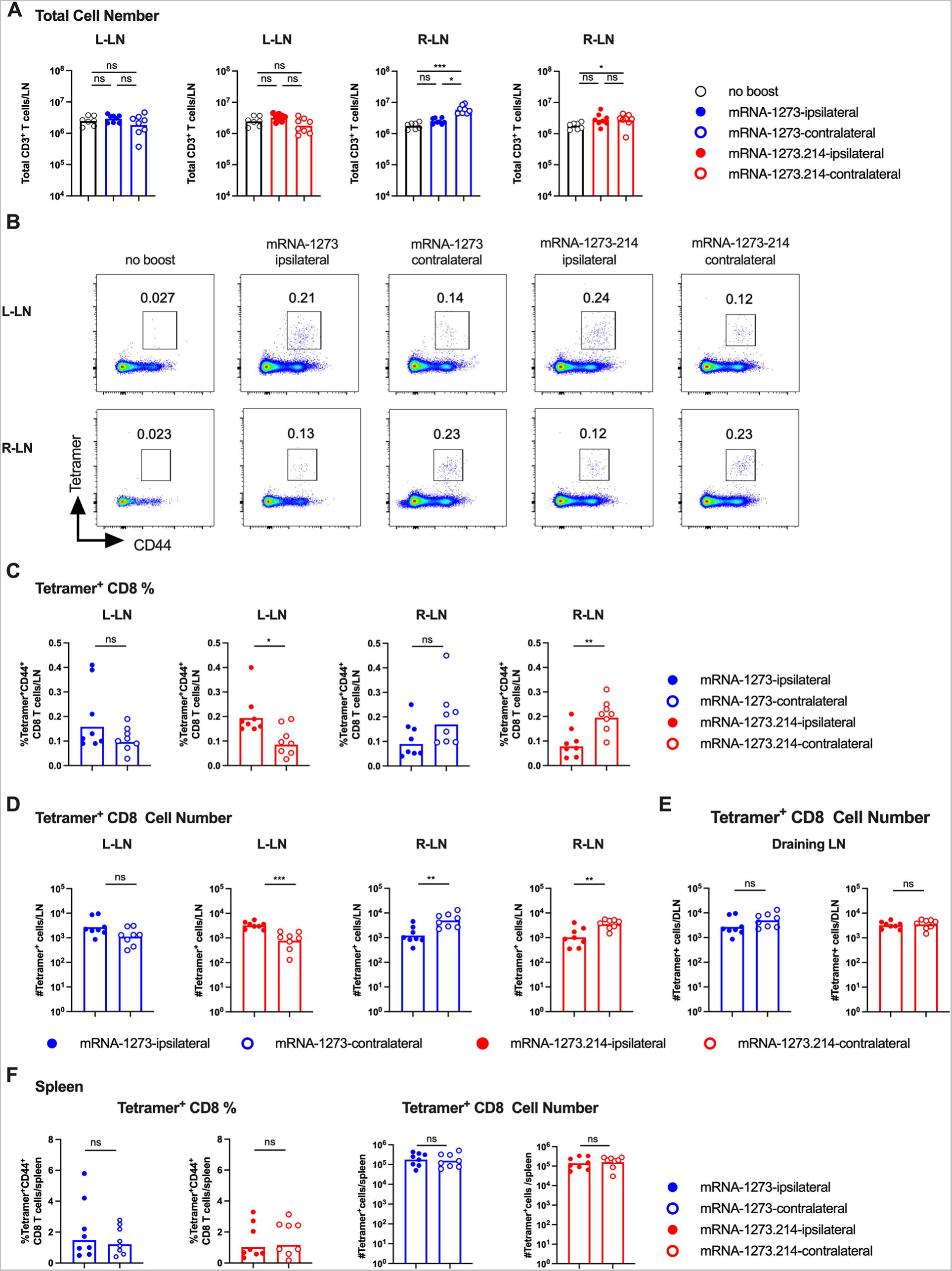
CD8^+^ T cell responses in the lymph nodes and spleen following boosting with mRNA-1273 or mRNA1273.214. Seven- to nine-week-old female K18-hACE2 mice were immunized with a primary two-dose vaccination series spaced three weeks apart in the left hind leg with mRNA-1273. Animals then were boosted 11 to 12 weeks later in the left (ipsilateral) or right (contralateral) leg with 1 µg of mRNA-1273 or mRNA-1273.214. Seven days after boosting, inguinal LNs from the left and right side as well as spleen were analyzed for CD8^+^ T cell responses by flow cytometry. **A**. Quantification of total number of CD3^+^ T cells in respective LNs. **B**. Representative flow cytometry scatter plots of spike-specific tetramer^+^CD44^+^CD8^+^ T cells in respective LNs. **C-D**. Spike-specific tetramer^+^CD44^+^CD8^+^ T cells in respective LNs: **C**, frequency, **D**, total cell number. **E.** Comparison of numbers of spike-specific tetramer^+^CD44^+^CD8^+^ T cells in the DLNs. **F**. Frequency and total cell number of spike-specific tetramer^+^CD44^+^CD8^+^ T cells in the spleen. Data are from two experiments (n = 7-8, each data point represents an individual mouse, column heights indicate geometric mean values. Statistical analyses: **A,** One-way ANOVA with Tukey’s post-test; **C, D, E, F,** unpaired two-tailed Mann-Whitney test: ns, not significant; * p < 0.05, ∗∗ p < 0.01, ∗∗∗ p < 0.001.

Follicular T helper cells (T_FH_: CD4^+^CD45^+^CXCR5^+^PD-1^+^BCL6^+^) shape B cell responses by instructing GC reactions and the process of somatic hypermutation and affinity selection ^33^. We assessed the impact of ipsilateral and contralateral boosting on T_FH_ cell responses in the LNs at 7 days post boosting (**Fig S6A-D**). Like that observed with CD8^+^ T cell responses, ipsilateral and contralateral leg boosting with mRNA-1273 or mRNA-214 vaccines elicited higher frequencies (**Fig S6A-B**) and numbers (**Fig S6C**, 3 to 10-fold, p < 0.05) of T_FH_ cells in the respective DLN, with reduced responses observed in the LN on the side opposite of boosting (**Fig S6B-C**). Ipsilateral and contralateral boosting elicited equivalent numbers of T_FH_ cells in the DLNs regardless of the vaccine used (**Fig S6D**). Likewise, ipsilateral and contralateral boosting with mRNA-1273 or mRNA-1273.214 mRNA elicited comparable frequencies and numbers of T_FH_ T cells in the spleen (**Fig S6E**).

### Effect of ipsilateral or contralateral boosting on protection against BA.1 challenge

To evaluate the protective activity of the mRNA vaccines following ipsilateral or contralateral boosting, the cohorts of K18-hACE2 mice were challenged nine weeks later via intranasal route with 10^4^ focus-forming units (FFU) of BA.1 (**Fig 6A**). One limitation of these studies is that BA.1 is inherently less pathogenic in rodents ^34–37^ and replicates to lower levels in both the upper and lower respiratory tracts of K18-hACE2 mice without the development of clinical signs of disease compared to earlier variants ^16,34^; as such, we focused our analysis on virological endpoints. Nasal turbinates and lungs were harvested at day 4 after BA.1 infection and assayed for viral RNA levels by qRT-PCR (**Fig 6B-C**). In the nasal turbinates of control mRNA vaccine immunized mice, although some variability was observed, moderate amounts of BA.1 viral RNA were detected (approximately 5 x 10^4^ copies of *N* transcript per mg) (**Fig 6B**). Ipsilateral and contralateral boosting with mRNA-1273 or mRNA-1273.214 vaccines conferred protection in the nasal turbinates, with 100-1,000-fold reductions (p < 0.05) in viral RNA compared to the control vaccinated mice. In the lungs of control vaccinated K18-hACE2 mice, 10^5^ to 10^7^ copies viral RNA per mg of tissue were detected (**Fig 6C**). Ipsilateral and contralateral boosting with either mRNA-1273 or mRNA-1273.214 substantially reduced (mRNA-1273, ipsilateral or contralateral, 90 to 190-fold; mRNA-1273.214, ipsilateral or contralateral 370- to 1,000-fold, p < 0.0001) levels of viral RNA in the lung compared to the control vaccinated mice with no statistically significant differences conferred by the site of boosting (p > 0.05). Consistent with these data, no differences in infectious BA.1 were detected by plaque assay after ipsilateral or contralateral boosting with mRNA-1273 or mRNA-1273.214, with each vaccine and immunization site resulting in significant reductions (14 to 26-fold, p < 0.0001) compared to control mRNA vaccinated and challenged controls (**Fig 6D**).

**Figure 6.**
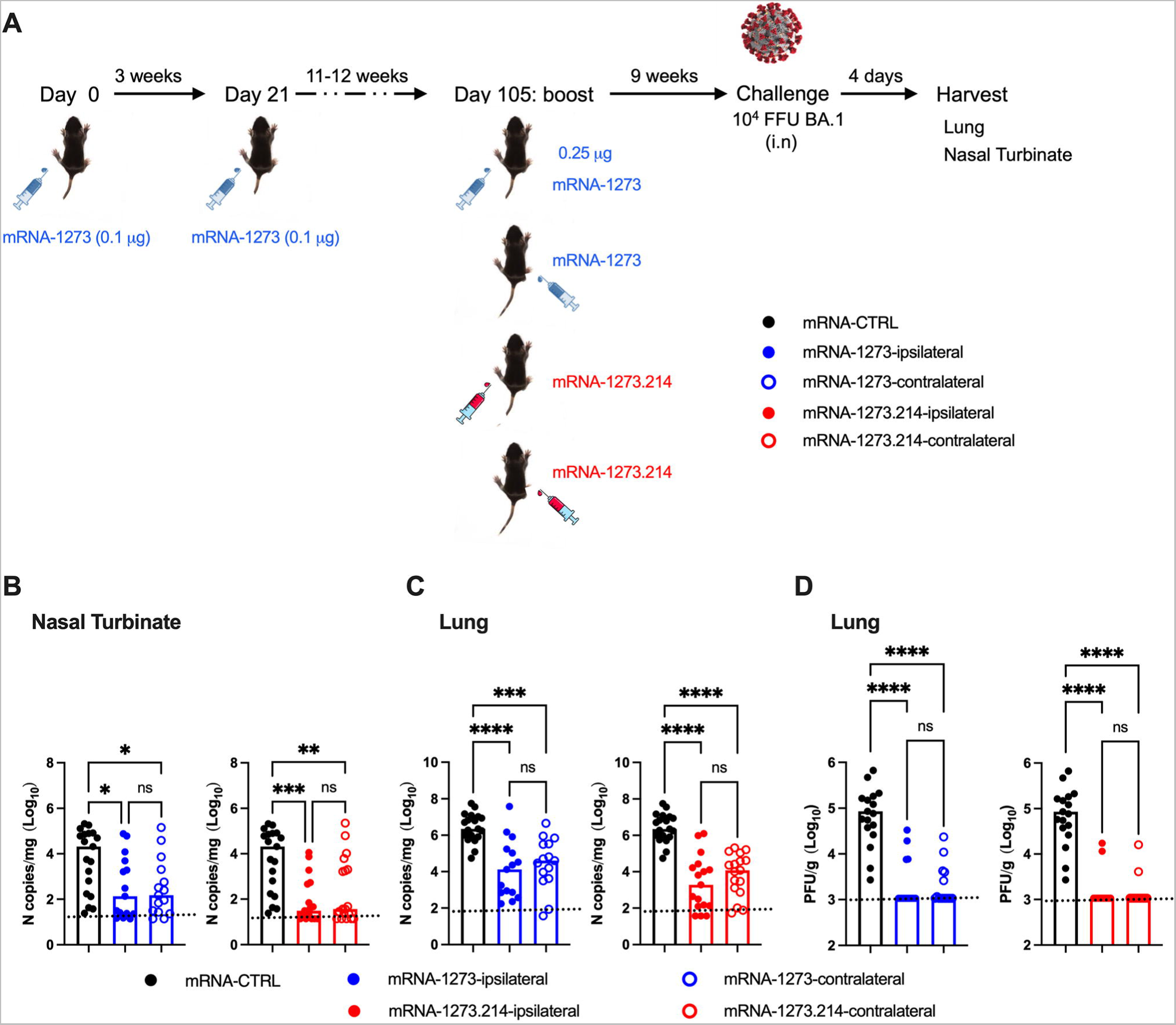
Protection against BA.1 infection in K18-hACE2 mice following boosting with mRNA-1273 or mRNA-1273.214. **A**. Scheme of immunizations and virus challenge. Seven- to nine-week-old female K18-hACE2 mice were immunized with a primary two-dose vaccination series spaced three weeks apart in the left hind leg with 0.1 μg of mRNA-1273. Animals were boosted 11 to 12 weeks later in the left (ipsilateral) or right (contralateral) leg with 0.25 μg of mRNA-1273 or bivalent mRNA-1273.214. Mice immunized with mRNA-control (left hind leg) were used as negative control group. Nine weeks after boosting, mice were challenged via intranasal route with 10^4^ focus-forming units (FFU) of SARS-CoV-2 BA.1. Viral RNA levels were determined at 4 dpi in the nasal turbinates (**B**) and lungs (**C**). **D**. Infectious virus in the lungs. Data are from two independent experiments (n = 15-18, each data point represents an individual mouse, column heights indicate median values. Statistical analyses: One-way ANOVA with Tukey’s post-test: ns, not significant; * p < 0.05, ∗∗ p < 0.01, ∗∗∗ p < 0.001, ∗∗∗∗ p < 0.0001.

## DISCUSSION

One of the key immunization strategies to mitigate waning immunity and reduced efficacy against SARS-CoV-2 variants that escape neutralization is boosting with vaccines that encode variant spike proteins. A frequent question regarding the clinical practice of boosting is which arm should be used, and whether boosting in the alternating arm impacts immunogenicity and protection against SARS-CoV-2. In this study, we evaluated in mice the impact of ipsilateral or contralateral site boosting of mRNA vaccines on the adaptive immune responses and protection against SARS-CoV2 challenge. We observed that ipsilateral and contralateral boosting elicited comparable overall levels of serum spike specific antibody as well as antigen-specific B and T cells responses. The differential impact of boosting site on protective response in the context of virus challenge was limited, as ipsilateral and contralateral boosting with mRNA-1273 or mRNA-1273.214 vaccines both conferred similar levels of virological protection against BA.1 infection.

Our data showed that ipsilateral and contralateral boosting elicited comparable levels of anti-spike IgG in serum at 4 weeks after immunization. Functional analysis of antibody responses revealed similar neutralizing activities against SARS-CoV-2 WA1/2020 D614G and BA.1 viruses with small differences that did not achieve statistical significance between booster site regimens. Our results in K18-hACE2 mice with COVID-19 mRNA vaccines generally are consistent with a two-dose vaccination study in conventional C57BL/6 mice using homologous influenza hemagglutinin (H1-HA) antigen, which showed comparable levels of serum anti-HA IgG at 8 days post boost, although that study did not examine neutralizing antibody activities ^21^. Our results contrast with this study when analyzing the effects of a heterologous antigen boost. In the influenza vaccine study, ipsilateral, but not contralateral, boosting with heterologous H3-HA enhanced H1-binding serum IgG levels, whereas contralateral boosting with H3-HA did not. While we did not observe such effects when comparing ipsilateral and contralateral administration of monovalent (mRNA-1273) and bivalent (mRNA-1273.214) mRNA vaccine boosters, the bivalent vaccine contains the original mRNA-1273, which likely affects recall responses against the original Wuhan-1 spike antigen. Future studies with a fully heterologous monovalent boost (*e.g*., mRNA-1273.529 (BA.1) or mRNA-1273.815 (XBB.1.5)) may allow for more direct comparisons to the influenza immunization experiments.

Our serum antibody data showed some similarities and differences with a recent study in C57BL/6 mice that used a two-dose homologous BNT162b2 (Pfizer-BioNTech) mRNA vaccination regimen ^24^. That study showed that ipsilateral boosting with BNT162b2 mRNA vaccine produced higher titers of serum antibodies with enhanced affinity against RBD of Wuhan-1 and Omicron strains at early (day 9) but not the late (week 19) timepoints post boost. However, like our results, no difference was observed in overall serum IgG levels after ipsilateral or contralateral boosting when analysis was performed with the spike proteins at either the early or later timepoints. Beyond pre-clinical studies in small animals, the data on the impact of ipsilateral versus contralateral boosting on antibody responses in human remain inconclusive. Two observational studies conducted among SARS-CoV-2 uninfected individuals who received the BNT162b2 mRNA vaccine reached different conclusions. Ziegler *et al* reported that spike-specific IgG levels did not differ between two groups that received a second dose of BNT162b2 on the ipsilateral (n = 147) or contralateral side (n = 156), although inhibitory activity was higher in the group boosted on the ipsilateral side at two weeks after the second dose based on an assay that measured inhibition of RBD binding to hACE2 ^22^. In contrast, Fazli *et al* reported that contralateral boosting (n = 440) with BNT162b2 mRNA vaccine was associated with slightly higher spike-and RBD-specific serum IgG titers compared to ipsilateral boosting (n = 507), and this effect increased over time from a 1.1-fold to a 1.4-fold effect by 14 months. Contralateral boosting also resulted in increases (1.3-fold to 4.0-fold) in neutralizing antibody titers against B.1.1.529 at 8- and 14-months, with lesser effects against the WA1/2020 D614G strain (no difference at 8-months and 2-fold difference at 14 months) ^23^.

To investigate the mechanisms by which boosting site selection might influence humoral immunity, we interrogated B cells in the LNs and spleen. Ipsilateral and contralateral leg boosting with mRNA-1273 or mRNA-1273.214 vaccines elicited robust GCB responses in the DLNs against Wuhan-1 and BA.1 spike proteins at 7 days post boost, with reduced responses observed in the LNs on the site opposite of boosting. As expected, bivalent mRNA-1273.214 elicited higher numbers of GCBs that bound BA.1 spike than mRNA-1273. Whereas contralateral boosting extended the GCB response to the contralateral DLN, ipsilateral or contralateral boosting with mRNA-1273 or mRNA-1273.214 vaccines overall induced similar levels of GCBs that bound Wuhan-1 or BA.1 spike in the DLNs. Our results are consistent with fate-mapping studies in mice, which showed that recruitment of antigen-experienced memory B cells to secondary GCs is inefficient ^38,39^, as the progeny of primed GC B cells account for only a subset of total GCBs. Our analysis on PBs/PCs responses revealed that ipsilateral and contralateral boosts elicited equivalent PBs/PCs responses in the respective DLNs and spleen, which is consistent with the similar spike- and RBD-specific IgG responses in serum by the two booster regimens. However, our results both agree and contrast with the two-dose influenza immunization study in mice ^21^, which showed that ipsilateral and contralateral boosting with homologous influenza hemagglutinin H1 antigen elicited comparable total GCB but better quality of H1 reactive GCBs in the DLN; in that study, ipsilateral boosting with the heterologous H3-HA antigen increased the breadth of GCBs that bound to H1-HA. Our GCB results also differ from the study with two homologous doses of BNT162b2 mRNA in mice ^24^, which showed that ipsilateral boosting induced higher number of GCBs that bound RBD of the Wuhan-1 strain in the ipsilateral DLN than that contralateral boosting elicited in the contralateral DLN.

Despite increasing data on antibody and B cell responses induced after immunization at different boosting sites in mice and humans, the results remain inconsistent and challenging to develop paradigms. Multiple factors in study design likely contribute to discrepant results among studies. (a) Vaccine platforms could impact responses. Whereas the influenza study used HA protein-based vaccines to interrogate the impact of ipsilateral or contralateral boosting, we used monovalent and bivalent mRNA vaccines. Vaccine platforms can affect the extent of GC responses in the DLN, as mRNA vaccines can induce persistent GC responses that last months compared to traditional protein-based platforms ^9,40–43^. (b) The frequency of boosters and interval between primary immunization could change the dynamics of B cell and antibody responses generated from ipsilateral and contralateral boosting. Whereas the previous studies in mice boosted after one-dose prime immunization, our study evaluated the impact of boost location following a two-dose primary vaccination series. Indeed, a third COVID-19 mRNA vaccine dose in humans has been associated with an increase in RBD-specific memory B cells both from expanded clones present after the second dose and the emergence of new clones ^44,45^. The boosting interval also differs between our study (12 weeks) and the prior BNT162b2 mRNA vaccine study (3 weeks) in mice ^24^, which has been shown to independently impact antibody responses ^46,47^. (c) The relative dosing of the booster could impact responses. Higher doses could result in prolonged antigen retention in the follicular dendritic cells in the DLN that differentially shape B cell and antibody responses ^48,49^. (d) The timing of assessment. The two human studies on ipsilateral and contralateral boosting may have reached different conclusions because they evaluated antibody responses at different time points, ranging from two weeks to 14 months ^22,23^. Analogously, the study in C57BL/6 mice with two homologous doses of BNT162b2 mRNA vaccine showed different affinity measurements at early and late time points after ipsilateral and contralateral boosting ^24^. Together, these data suggest that at the early phase after boosting, memory B cells in the DLN on the ipsilateral side with respect to priming likely differentiate into PB/PC rapidly after ipsilateral boosting, resulting in earlier serum antibody production than contralateral boosting. In comparison, after contralateral boosting, in the contralateral DLN, newly recruited antigen-specific B cells from the naive pool expand, affinity mature, and differentiate into plasmabasts to produce antibodies. (e) Antigenic distance between primary and boost vaccines could impact the responses. As shown in the influenza study ^21^, ipsilateral boosting with a heterologous antigen increased cross reactive-antibody responses more than contralateral boosting, which might be better at overcoming immune imprinting. The bivalent vaccine in our study contains the original mRNA-1273, which likely affects recall responses against the original Wuhan-1 spike antigen. For mRNA vaccines, more study of the effects of ipsilateral and contralateral boosting is likely needed for evaluating fully heterologous monovalent vaccines.

T cell immune responses contribute to protection against SARS-CoV-2 infection and disease, especially in the setting of poor neutralizing antibody responses against variants ^8,29–32^. We interrogated spike-specific CD8 and total T_FH_ cell responses locally and systemically. Ipsilateral and contralateral boosting with mRNA-1273 or mRNA-1273.214 vaccines both elicited enhanced spike-specific CD8^+^ T cell and overall T_FH_ responses in the spleen and DLN at 7 days post boosting, with reduced responses observed in LN at the site opposite of boosting. Notwithstanding these data, ipsilateral and contralateral boosting with mRNA-1273 or mRNA- 1273.214 vaccines elicited comparable spike-specific CD8^+^ T cells and overall T_FH_ responses in the spleen and respective DLN. These results are consistent with a study in C57BL/6 mice that used self-amplifying mRNA vaccine encoding the influenza A virus nucleoprotein ^50^ and showed comparable antigen-specific CD4^+^ and CD8^+^ T cells in the spleen 28 days after ipsilateral and contralateral boosting. Our results with T_FH_ cells differ from this study, as we characterized these cells in the spleen and respective DLNs at 7 days after ipsilateral and contralateral boosting. Our CD8^+^ T cell results also differ from a study in humans, which showed that frequencies of spike-specific CD8^+^ T cell in the blood were higher after ipsilateral than contralateral boosting ^22^. The reasons for the differences are not clear beyond the different anatomic compartments (spleen versus blood) sampled.

We uniquely assessed the impact of ipsilateral and contralateral boosting on vaccine-mediated protection against virus challenge. Boosting with either mRNA-1273 or mRNA-1273.214 efficiently reduced BA.1 viral RNA levels in the upper and lower respiratory tracts with a trend toward higher level of protection in animals boosted with mRNA-1273.214, as observed previously ^16^,. Notably, our data showed that the site of boosting with mRNA-1273 or mRNA-1273.214 vaccines did not affect protection against BA.1 as judged by measurement of viral RNA and infectious virus in the nasal turbinates and lungs. Boosting at either site with mRNA-1273 or mRNA-1273.214 reduced the levels of viral infection, with each vaccine and immunization site resulting in significant reductions compared to unvaccinated, challenged controls. Overall, these data in mice show that the limited differences in overall immune response induced by ipsilateral or contralateral boosting with prototype (mRNA-1273) or variant-matched bivalent (mRNA-1273.214) do not substantively impact virological protection against challenge by a SARS-CoV-2 Omicron strain.

We acknowledge several limitations in our study. (a) We evaluated humoral or cellular immune responses at only one time point following boosting. Longitudinal studies after ipsilateral and contralateral boosting are needed to assess whether the dynamics of immune responses change over time. (b) Our study evaluated immune responses after one ipsilateral or contralateral boost after a primary vaccination series. How additional boosting on the contralateral side shifts the recall responses remains to be determined. (c) We used monovalent and bivalent mRNA vaccine formulations in our boosting schemes in part because of when the studies were initiated. Experiments that test monovalent heterologous boosting with more contemporary vaccines (encoding spikes from BA.1, BA.5, XBB.1.5, or emerging strains) are needed to determine how the boosting site affects or potentially overcomes immune imprinting. (d) We performed studies only in uninfected mice receiving vaccines. Future studies on boost location are needed in the setting of antecedent SARS-CoV-2 infection to extrapolate our findings to conditions of hybrid immunity. (e) We did not analyze the impact of boost location on bone marrow-derived long-lived plasma cells (LLPCs). LLPCs constitutively secrete high levels of antibodies for extended periods of time and dominantly contribute to serum antibody levels at times remote from vaccination or infection. (f) Finally, studies with different vaccine platforms as well as other animal models and ultimately, humans are required for corroboration.

Our studies in K18-hACE2 mice provide evidence that ipsilateral and contralateral boosting with mRNA vaccines elicit comparable immune responses, and this was associated with equivalent control of infection by a SARS-CoV-2 Omicron challenge strain. Thus, at least in mice, the boost site location for mRNA vaccines appears to have limited impacts on immunogenicity and protection against SARS-CoV-2.

## Acknowledgements

This study was supported by the National Institutes of Health (NIAID Centers of Excellence for Influenza Research and Response (CEIRR) contracts 75N93021C00014 and 75N93019C00051, to M.S.D.) and a sponsored Research Agreement with Moderna.

## Author contributions

B.Y. performed serum antibody and viral burden analysis. B.Y. and S.M.S. performed experiments in mice. C.Y.L. and B.Y. performed B cell staining and analysis. P.D. performed T cell staining and analysis. S.M.E. and D.K.E. provided mRNA vaccines. L.B.T. and M.S.D. designed studies and supervised the research. B.Y. and M.S.D. wrote the initial draft, with all other authors providing editorial comments.

## SUPPLEMENTAL FIGURE LEGENDS

**Figure S1. Serum neutralization of WA1/2020 D614G and BA.1 viruses, Related to** Figure 2. Seven- to nine-week-old female K18-hACE2 transgenic mice were immunized with a primary mRNA-1273 vaccination series and then boosted in the ipsilateral or contralateral leg with mRNA-1273 or mRNA-1273.214. Neutralizing antibody responses against WA1/2020 D614G and BA.1 were assessed from serum samples from one day before (**A**) or four weeks after (**B**) booster dose vaccines (n = 15-18). Neutralization curves corresponding to individual mice are shown for the indicated vaccines. Each point represents the mean of two technical replicates.

**Figure S2. Flow cytometry gating strategies for B cells in the LN and spleen, Related to** Figures 3**, 4, S3, and S4**. **A.** GCBs were gated for lymphocytes (FSC-A/SSC-A), singlets (FSC-H/FSC-A), live cells (Viability dye eF506^-^), CD19^+^, IgD ^low^, Fas^+^GL7^+^, decoy Oval^-^ (non-specific binding negative), followed by Wuhan-1 spike (BV421^+^BV786^+^) or BA.1 spike (APC^+^AF594^+^). **B.** PBs/PCs were gated for lymphocytes (FSC-A/SSC-A), singlets (FSC-H/FSC-A), live cells (Viability dye eF506^-^), CD19^+^, IgD ^low^, CD138^+^TACI^+^, decoy Oval- (non-specific binding negative), followed by Wuhan-1-spike (BV421^+^BV786^+^) or BA.1 spike (APC^+^AF594^+^).

**Figure S3. Frequency of germinal center B cells and plasmablast/plasma cells in the lymph nodes following boosting with mRNA-1273 or mRNA1273.214, Related to** Figures 3 **and 4**. (**A-C**) GCBs. Frequency of all GCBs (**A**), Wuhan-1 spike-reactive GCBs (**B**), and BA.1-spike reactive (**C**) GCBs among antigen-experienced B cells (CD19^+^IgD^lo^). (**D-F**) PB/PCs. Frequency of all PB/PCs (**D**), Wuhan-1 spike-reactive PB/PCs (**E**), and BA.1-spike reactive (**F**) PB/PCs among antigen-experienced B cells (CD19^+^IgD^lo^). Data are from two independent experiments (n = 7-8, each data point represents an individual mouse, column heights indicate median values). Statistical analyses: **A, D,** One-way ANOVA with Tukey’s post-test; **B-C, E-F,** unpaired two-tailed Mann-Whitney test: ns, not significant; ∗ p < 0.05, ∗∗ p < 0.01, ∗∗∗ p < 0.001, ∗∗∗∗ p < 0.0001.

**Figure S4. Plasmablast/plasma cell responses in the spleen following boosting with mRNA-1273 or mRNA1273.214.** Seven- to nine-week-old female K18-hACE2 transgenic mice were immunized with a primary mRNA-1273 vaccination series and then boosted 11 to 12 weeks later in the ipsilateral or contralateral leg with mRNA-1273 or mRNA-1273.214. Seven days after boosting, spleens were harvested and analyzed for PBs/PCs responses by flow cytometry. **A**. Total number of CD19^+^IgD^lo^CD138^+^TACI^+^PB/PC cells. **B**. Representative flow cytometry scatter plots (left) and quantification (right) of Wuhan-1 spike-reactive PBs/PCs. **C**. Representative flow cytometry scatter plots (left) and quantification (right) of BA.1 spike reactive PBs/PCs. Data are from two independent experiments (n = 7-8, each data point represents an individual mouse, column heights indicate geometric mean values). Statistical analyses: **A,** One-way ANOVA with Tukey’s post-test; **B-C,** unpaired two-tailed Mann-Whitney test: ns, not significant; ∗∗∗∗ p < 0.0001.

**Figure S5. Gating strategies for analyzing spike specific CD8^+^ T and total T_FH_ cells in the LN and spleen, Related to** Figures 5**, S6.** Cells were gated for lymphocytes (FSC-A/SSC-A), singlets (FSC-H/FSC-A; SSC-H/SSC-A), CD45^+^, CD3^+^, CD8^+^, CD4^+^, followed by tetramer^+^CD44+ (tetramer^+^CD8^+)^ and PD-1^+^CXCR5^+^ (T_FH_).

**Figure S6. T_FH_ cell responses in the lymph node and spleen following boosting with mRNA-1273 or mRNA1273.214.** Seven- to nine-week-old female K18-hACE2 mice were immunized with a primary two-dose vaccination series spaced three weeks apart in the left hind leg with mRNA-1273. Animals then were boosted 11 to 12 weeks later in the left (ipsilateral) or right (contralateral) leg with 1 µg of mRNA-1273 or mRNA-1273.214. Seven days after boosting, inguinal LNs from the left, right side and spleens were analyzed for T_FH_ responses by flow cytometry. **A**. Representative flow cytometry scatter plots of PD1^+^CXCR5^+^ T_FH_ cells in the LN. **B**. Frequency PD1^+^CXCR5^+^ T_FH_ cells in respective LNs. **C.** Total cell number of PD1^+^CXCR5^+^ T_FH_ cells in the DLNs. **D.** Comparison of numbers of T_FH_ cells in the DLNs. **E**. Frequency and total cell number of T_FH_ cells in the spleen. Data are from two experiments (n = 7-8, each data point represents an individual mouse, column heights indicate geometric mean values). Statistical analyses: **B**, **C, D, E,** unpaired two-tailed Mann-Whitney test: ns, not significant; * p < 0.05, ∗∗ p < 0.01, ∗∗∗ p < 0.001.

## METHODS

### Cells

African green monkey Vero-TMPRSS2 ^51^ and Vero-hACE2-TMPRRS2 ^52^ cells were cultured at 37°C in Dulbecco’s Modified Eagle medium (DMEM) supplemented with 10% fetal bovine serum (FBS), 10 mM HEPES pH 7.3, 1 mM sodium pyruvate, 1× non-essential amino acids, and 100 U/mL of penicillin–streptomycin. Vero-TMPRSS2 cells were cultured in medium supplemented with 5 μg/mL of blasticidin. Vero-hACE2-TMPRSS2 cells were propagated in medium supplemented with 5 μg/mL of blasticidin and 10 µg/mL of puromycin. All cells were routinely tested negative for mycoplasma using a PCR-based assay.

### Viruses

The WA1/2020 strain with a D614G substitution was described previously ^53^. The BA.1 (B.1.1.529) isolate (hCoV-19/USA/WI-WSLH-221686/2021) was passaged once on Vero-TMPRSS2 cells and described previously ^54^. All viruses were subjected to next-generation sequencing to confirm substitutions. All virus experiments were performed with Institutional Biosafety Committee approval in an approved biosafety level 3 (BSL-3 or A-BSL3) facility at Washington University School of Medicine.

### Mice

Heterozygous K18-hACE2 C57BL/6J mice (strain: 2B6.Cg-Tg(K18-ACE2)2Prlmn/J, Cat # 034860) were obtained from The Jackson Laboratory. Animal studies were carried out in accordance with the recommendations in the Guide for the Care and Use of Laboratory Animals of the National Institutes of Health. The protocols were approved by the Institutional Animal Care and Use Committee at the Washington University School of Medicine (assurance number A3381–01). Virus inoculations were performed under anesthesia that was induced and maintained with ketamine hydrochloride and xylazine, and all efforts were made to minimize animal suffering. Experiments were neither randomized nor blinded.

### mRNA vaccine and lipid nanoparticle production process

Sequence-optimized mRNA encoding prefusion-stabilized Wuhan-1 (mRNA-1273) and BA.1 (mRNA-1273.529, the Omicron gene component in mRNA-1273.214) SARS-CoV-2 S-2P proteins were synthesized *in vitro* using an optimized T7 RNA polymerase-mediated transcription reaction with complete replacement of uridine by N1m-pseudouridine ^55^. Bivalent mRNA-1273.214 vaccine is a 1:1 bench side mix of separately formulated mRNA-1273 and mRNA-1273.529 vaccines.

A non-translating control mRNA was synthesized and formulated into lipid nanoparticles as previously described ^56^. The reaction included a DNA template containing the immunogen open-reading frame flanked by 5’ untranslated region (UTR) and 3’ UTR sequences and was terminated by an encoded poly A tail. After RNA transcription, the cap-1 structure was added using the vaccinia virus capping enzyme and 2L-*O*-methyltransferase (New England Biolabs). The mRNA was purified, sterile filtered, and kept frozen at –20°C until further use.

The mRNA was encapsulated in a lipid nanoparticle as described previously ^57^. Vials were filled with formulated lipid nanoparticle and stored frozen at –20°C until further use. The vaccine product underwent analytical characterization, which included the determination of particle size and polydispersity, encapsulation, mRNA purity, double-stranded RNA content, osmolality, pH, endotoxin, and bioburden, and the material was deemed acceptable for *in vivo* study.

### Viral antigens

Recombinant soluble spike and RBD proteins from Wuhan-1 and BA.1 SARS-CoV-2 strains were expressed as described ^58,59^ Recombinant proteins were produced in Expi293F cells (ThermoFisher) by transfection of DNA using the ExpiFectamine 293 Transfection Kit (ThermoFisher). Supernatants were harvested 3 days post-transfection, and recombinant proteins were purified using Ni-NTA agarose (ThermoFisher), then buffer exchanged into PBS and concentrated using Amicon Ultracel centrifugal filters (EMD Millipore).

### ELISA

Purified recombinant Wuhan-1 or BA.1 spike or RBD proteins were coated onto 96-well Maxisorp clear plates at 2 μg/mL (spike) or 4 μg/mL (RBD) in 50 mM Na_2_CO_3_ pH 9.6 (50 μL) overnight at 4°C. Coating buffers were aspirated, and wells were blocked with 200 μL of 1X PBS + 0.05% Tween-20 + 2% BSA + 0.02% NaN_3_ (Blocking buffer, PBSTBA) overnight at 4°C. Sera were serially diluted in blocking buffer and added to the plates. Plates were incubated for 1 h at room temperature and then washed 3 times with PBST, followed by addition of 50 μL of 1:2000 anti-mouse IgG-HRP (Southern Biotech Cat. #1030-05) in PBST. Following a 1 h incubation at room temperature, plates were washed 3 times with PBST and 50 μL of 1-Step Ultra TMB-ELISA was added (ThermoFisher Cat. #34028). Following a 2 to 5-min incubation, reactions were stopped with 50 μL of 2 M H_2_SO_4_. The absorbance of each well at 450 nm was determined using a microplate reader (BioTek) within 5 min of addition of sulfuric acid. The endpoint serum dilution was calculated with curve fit analysis of optical density (OD) values for serially diluted sera with a cut-off value set to six times the mean of the background signal.

### Focus reduction neutralization test

Serial dilutions of sera were incubated with 10^2^ focus-forming units (FFU) of WA1/2020 D614G or BA.1 for 1 h at 37°C. Antibody-virus complexes were added to Vero-TMPRSS2 cell monolayers in 96-well plates and incubated at 37°C for 1 h. Subsequently, cells were overlaid with 1% (w/v) methylcellulose in MEM. Plates were harvested 30 h (WA1/2020 D614G) or 70 h (BA.1) later by removing overlays and fixed with 4% PFA in PBS for 20 min at room temperature. Plates were washed and sequentially incubated with an oligoclonal pool (SARS2-02, -08, -09, -10, -11, -13, -14, -17, -20, -26, -27, -28, -31, -38, -41, -42, -44, -49,, -57, -62, -64, -65, -67, and -71 ^60^ of anti-spike murine antibodies (including cross-reactive mAbs to SARS-CoV) and HRP-conjugated goat anti-mouse IgG (Sigma Cat # A8924, RRID: AB_258426) in PBS supplemented with 0.1% saponin and 0.1% bovine serum albumin. SARS-CoV-2-infected cell foci were visualized using TrueBlue peroxidase substrate (KPL) and quantitated on an ImmunoSpot microanalyzer (Cellular Technologies).

### Mouse experiments

Seven- to nine-week-old female K18-hACE2 C57BL/6 mice were immunized three weeks apart with 0.1 μg of mRNA-1273 vaccine in 50 µl of PBS via intramuscular injection in the left hind leg. Animals were boosted 12 weeks later in the left (ipsilateral) or right (contralateral) leg with 0.25 µg of mRNA-1273 or bivalent mRNA-1273.214 vaccine (1:1 mixture of mRNA-1273 and the BA.1-matched mRNA-1273.529). Animals were bled one day before and four weeks after boosting for antibody analysis.

To evaluate the protective activity of the mRNA vaccines nine weeks after boosting, the cohorts of K18-hACE2 mice were challenged via intranasal route with 10^4^ focus-forming units (FFU) of BA.1. Animals were euthanized at 4 dpi. Nasal turbinates and lung were harvested for virological and immunological analyses. For B and T cells analysis, the same vaccination scheme was applied but animals were boosted with 1 µg of mRNA-1273 or bivalent mRNA-1273.214 in the ipsilateral (left) or contralateral (right) leg. Seven days after boosting, the left and right inguinal LN and spleen were harvested for immune cells analysis.

### SARS-CoV-2 spike probe generation

Recombinant spike proteins were biotinylated with EZ-Link™ NHS-PEG4-Biotin (ThermoFisher, Cat. # A39259) for 2 h at 4°C and processed through Zeba spin desalting columns (ThermoFisher) to remove excess unbound biotin, purified biotinylated full-length Wuhan-1 and BA.1 spike proteins were subsequently fluorescently conjugated by mixing with fluorophore labelled streptavidin at 1:1.5 molar ratio for 1 h at 4°C (Wuhan-1 spike: strepavidin-BV421 or -BV786; BA.1-spike: streptavidin-APC or -AF594) prior to antigen specific B cell staining.

### B cell phenotyping

Seven days after boosting, ipsilateral and contralateral inguinal LNs and spleens were collected, and single cell suspensions were generated after tissue disruption and passage through a 70-μm cell strainer. Splenocytes were pelleted by centrifugation, and erythrocytes were lysed using ACK lysis buffer (Thermo Fisher). Cells were collected in DMEM with 10% FBS on ice. All staining steps were performed at 4°C in PBS with 2% heat inactivated FBS (FACS buffer). Single cell suspensions were blocked for FcγR binding with anti-CD16/CD32 monoclonal antibody (eBioscience, #14-0161-82) on ice for 15 min prior to staining. Cells were subsequently incubated for 60 min on ice with a pool of biotin-streptavidin conjugated recombinant spike protein (Wuhan-1-spike BV421, Wuhan-1-spike BV786, BA.1-spike APC, BA.1-spike AF594, oval-biotin-streptavidin-BUV737) in FACS buffer (2% FBS and 2 mM EDTA in PBS), washed twice, then stained for 30 min on ice with a cocktail of labeled mAbs including Fixable Viability dye eFluor506, CD4-AF700 (Biolegend, #100536), CD19-PerCP-Cy5.5. (BD, #551001), IgD-BV711 (BD, #564275), Fas-PE-Cy7 (BD, #557653), GL7-FITC (Biolegend, #144604), CD138-BV605 (Biolegend, #142516), TACI-PE (Biolegend, #133403). Cells were washed twice with FACS buffer, fixed with 1% paraformaldehyde (PFA) for 30 min prior to data acquisition. Data were acquired on an Aurora (Cytek) spectral flow cytometer and analyzed in FlowJo v10 software.

### T cell phenotyping

For T cell analysis, single-cell suspensions were incubated with FcγR antibody (clone 93, BioLegend) to block non-specific antibody binding, followed by staining with a cocktail of labeled monoclonal antibodies, including Fixable Viability dye eFluor506, CD45 (BUV395; Clone 30-F11; Cat: 564279; BD Biosciences), CD3 (BV711; Clone 145-2C11; Cat: 563123; BD Biosciences), CD8α (PerCP/Cy5.5; Clone 53-6.7; Cat: 100734; BioLegend), CD4 (BV785; Clone GK1.5; Cat: 100453; BioLegend), CD44 (APC/Cy7; Clone IM7; Cat: 103028; BioLegend), PD-1 PE/Cy7 (clone RMP1-30; Biolegend Cat# 109110), CXCR-5 BV421 (clone L138D7; Biolegend, Cat# 145511), and APC-labeled SARS-CoV-2 S-specific tetramer (MHC class I tetramer, residues 539–546, VNFNFNGL, H-2K^b^) for 60 min at room temperature. Cells were washed twice with FACS buffer and fixed with 2% paraformaldehyde for 5 min before data acquisition. Data were acquired on an Aurora (Cytek) spectral flow cytometer and analyzed with FlowJo v10 software.

### Measurement of viral RNA

Tissues were weighed and homogenized with zirconia beads in a MagNA Lyser instrument (Roche Life Science) in 1 ml of DMEM medium supplemented with 2% heat inactivated FBS. Tissue homogenates were clarified by centrifugation at 10,000 rpm for 5 min and stored at −80°C. RNA was extracted using the MagMax mirVana. Total RNA isolation kit (Thermo Fisher Scientific) on the Kingfisher Flex extraction robot (Thermo Fisher Scientific). RNA was reverse transcribed and amplified using the TaqMan RNA-to-CT 1-Step Kit (Thermo Fisher Scientific). Reverse transcription was carried out at 48°C for 15 min followed by 2 min at 95°C. Amplification was accomplished over 50 cycles as follows: 95°C for 15 s and 60°C for 1 min. Copies of SARS-CoV-2 *N* gene RNA in samples were determined using a published assay ^61^.

### Viral plaque assays

Plaque assays for titration of infectious virus were performed on Vero-hACE2-hTRMPSS2 cells in 24-well plates. Lung tissue homogenates were serially diluted tenfold, starting at 1:10, in cell infection medium (DMEML+L2% FBSL+L100Lu/ml penicillin– streptomycin). Two hundred microliters of the diluted virus were added to a single well per dilution per sample. After 1 h at 37 °C, the cells were overlayed with one milliliter of 1% methylcellulose in MEM supplemented with 2% FBS was added. Ninety-six hours after virus inoculation, the cells were fixed with 10% formalin, and the monolayer was stained with crystal violet (0.05% (w/v) in 25% methanol in water) for 30 min at 20°C. Plaque numbers were counted and used to calculate the PFU per gram.

### Statistical analysis

Statistical significance was assigned when p values were < 0.05 using GraphPad Prism version 9.3. Tests, number of animals, median, mean or geometric mean values, and statistical comparison groups are indicated in the Figure legends. Changes in viral RNA levels, or serum antibody responses were compared to unvaccinated animals and were analyzed by one-way ANOVA with a multiple comparisons correction, Mann-Whitney test, or Wilcoxon signed-rank test depending on the type of results, number of comparisons, and distribution of the data.

## REFERENCES

1. Watson, O.J., Barnsley, G., Toor, J., Hogan, A.B., Winskill, P., and Ghani, A.C. (2022). Global impact of the first year of COVID-19 vaccination: a mathematical modelling study. Lancet Infect Dis 22, 1293–1302. 10.1016/S1473-3099(22)00320-6.

2. Cele, S., Jackson, L., Khoury, D.S., Khan, K., Moyo-Gwete, T., Tegally, H., San, J.E., Cromer, D., Scheepers, C., Amoako, D.G., et al. (2022). Omicron extensively but incompletely escapes Pfizer BNT162b2 neutralization. Nature 602, 654–656. 10.1038/s41586-021-04387-1.

3. Liu, L., Iketani, S., Guo, Y., Chan, J.F., Wang, M., Liu, L., Luo, Y., Chu, H., Huang, Y., Nair, M.S., et al. (2022). Striking antibody evasion manifested by the Omicron variant of SARS-CoV-2. Nature 602, 676–681. 10.1038/s41586-021-04388-0.

4. Viana, R., Moyo, S., Amoako, D.G., Tegally, H., Scheepers, C., Althaus, C.L., Anyaneji, U.J., Bester, P.A., Boni, M.F., Chand, M., et al. (2022). Rapid epidemic expansion of the SARS-CoV-2 Omicron variant in southern Africa. Nature 603, 679–686. 10.1038/s41586-022-04411-y.

5. Lau, J.J., Cheng, S.M.S., Leung, K., Lee, C.K., Hachim, A., Tsang, L.C.H., Yam, K.W.H., Chaothai, S., Kwan, K.K.H., Chai, Z.Y.H., et al. (2023). Real-world COVID-19 vaccine effectiveness against the Omicron BA.2 variant in a SARS-CoV-2 infection-naive population. Nat Med 29, 348–357. 10.1038/s41591-023-02219-5.

6. Goel, R.R., Painter, M.M., Apostolidis, S.A., Mathew, D., Meng, W., Rosenfeld, A.M., Lundgreen, K.A., Reynaldi, A., Khoury, D.S., Pattekar, A., et al. (2021). mRNA vaccines induce durable immune memory to SARS-CoV-2 and variants of concern. Science 374, abm0829. 10.1126/science.abm0829.

7. Kim, W., Zhou, J.Q., Horvath, S.C., Schmitz, A.J., Sturtz, A.J., Lei, T., Liu, Z., Kalaidina, E., Thapa, M., Alsoussi, W.B., et al. (2022). Germinal centre-driven maturation of B cell response to mRNA vaccination. Nature 604, 141–145. 10.1038/s41586-022-04527-1.

8. Sette, A., and Crotty, S. (2021). Adaptive immunity to SARS-CoV-2 and COVID-19. Cell 184, 861–880. 10.1016/j.cell.2021.01.007.

9. Turner, J.S., O’Halloran, J.A., Kalaidina, E., Kim, W., Schmitz, A.J., Zhou, J.Q., Lei, T., Thapa, M., Chen, R.E., Case, J.B., et al. (2021). SARS-CoV-2 mRNA vaccines induce persistent human germinal centre responses. Nature 596, 109–113. 10.1038/s41586-021-03738-2.

10. Collier, A.Y., Yu, J., McMahan, K., Liu, J., Chandrashekar, A., Maron, J.S., Atyeo, C., Martinez, D.R., Ansel, J.L., Aguayo, R., et al. (2021). Differential Kinetics of Immune Responses Elicited by Covid-19 Vaccines. N Engl J Med 385, 2010–2012. 10.1056/NEJMc2115596.

11. Pegu, A., O’Connell, S., Schmidt, S.D., O’Dell, S., Talana, C.A., Lai, L., Albert, J., Anderson, E., Bennett, H., Corbett, K.S., et al. (2021). Durability of mRNA-1273-induced antibodies against SARS-CoV-2 variants. bioRxiv. 10.1101/2021.05.13.444010.

12. Goldberg, Y., Mandel, M., Bar-On, Y.M., Bodenheimer, O., Freedman, L., Haas, E.J., Milo, R., Alroy-Preis, S., Ash, N., and Huppert, A. (2021). Waning Immunity after the BNT162b2 Vaccine in Israel. N Engl J Med 385, e85. 10.1056/NEJMoa2114228.

13. Menegale, F., Manica, M., Zardini, A., Guzzetta, G., Marziano, V., d’Andrea, V., Trentini, F., Ajelli, M., Poletti, P., and Merler, S. (2023). Evaluation of Waning of SARS-CoV-2 Vaccine-Induced Immunity: A Systematic Review and Meta-analysis. JAMA Netw Open 6, e2310650. 10.1001/jamanetworkopen.2023.10650.

14. Ferdinands, J.M., Rao, S., Dixon, B.E., Mitchell, P.K., DeSilva, M.B., Irving, S.A., Lewis, N., Natarajan, K., Stenehjem, E., Grannis, S.J., et al. (2022). Waning 2-Dose and 3-Dose Effectiveness of mRNA Vaccines Against COVID-19-Associated Emergency Department and Urgent Care Encounters and Hospitalizations Among Adults During Periods of Delta and Omicron Variant Predominance - VISION Network, 10 States, August 2021-January 2022. MMWR Morb Mortal Wkly Rep 71, 255–263. 10.15585/mmwr.mm7107e2.

15. Pajon, R., Doria-Rose, N.A., Shen, X., Schmidt, S.D., O’Dell, S., McDanal, C., Feng, W., Tong, J., Eaton, A., Maglinao, M., et al. (2022). SARS-CoV-2 Omicron Variant Neutralization after mRNA-1273 Booster Vaccination. N Engl J Med. 10.1056/NEJMc2119912.

16. Ying, B., Scheaffer, S.M., Whitener, B., Liang, C.Y., Dmytrenko, O., Mackin, S., Wu, K., Lee, D., Avena, L.E., Chong, Z., et al. (2022). Boosting with variant-matched or historical mRNA vaccines protects against Omicron infection in mice. Cell 185, 1572–1587 e1511. 10.1016/j.cell.2022.03.037.

17. Scheaffer, S.M., Lee, D., Whitener, B., Ying, B., Wu, K., Liang, C.Y., Jani, H., Martin, P., Amato, N.J., Avena, L.E., et al. (2023). Bivalent SARS-CoV-2 mRNA vaccines increase breadth of neutralization and protect against the BA.5 Omicron variant in mice. Nat Med 29, 247–257. 10.1038/s41591-022-02092-8.

18. Garcia-Beltran, W.F., St Denis, K.J., Hoelzemer, A., Lam, E.C., Nitido, A.D., Sheehan, M.L., Berrios, C., Ofoman, O., Chang, C.C., Hauser, B.M., et al. (2022). mRNA-based COVID-19 vaccine boosters induce neutralizing immunity against SARS-CoV-2 Omicron variant. Cell 185, 457–466 e454. 10.1016/j.cell.2021.12.033.

19. Wang, Q., Bowen, A., Valdez, R., Gherasim, C., Gordon, A., Liu, L., and Ho, D.D. (2023). Antibody Response to Omicron BA.4-BA.5 Bivalent Booster. N Engl J Med 388, 567–569. 10.1056/NEJMc2213907.

20. Wang, Q., Guo, Y., Bowen, A., Mellis, I.A., Valdez, R., Gherasim, C., Gordon, A., Liu, L., and Ho, D.D. (2024). XBB.1.5 monovalent mRNA vaccine booster elicits robust neutralizing antibodies against XBB subvariants and JN.1. Cell Host Microbe. 10.1016/j.chom.2024.01.014.

21. Kuraoka, M., Yeh, C.H., Bajic, G., Kotaki, R., Song, S., Windsor, I., Harrison, S.C., and Kelsoe, G. (2022). Recall of B cell memory depends on relative locations of prime and boost immunization. Sci Immunol 7, eabn5311. 10.1126/sciimmunol.abn5311.

22. Ziegler, L., Klemis, V., Schmidt, T., Schneitler, S., Baum, C., Neumann, J., Becker, S.L., Gartner, B.C., Sester, U., and Sester, M. (2023). Differences in SARS-CoV-2 specific humoral and cellular immune responses after contralateral and ipsilateral COVID-19 vaccination. EBioMedicine 95, 104743. 10.1016/j.ebiom.2023.104743.

23. Fazli, S., Thomas, A., Estrada, A.E., Ross, H.A., Xthona Lee, D., Kazmierczak, S., Slifka, M.K., Montefiori, D., Messer, W.B., and Curlin, M.E. (2024). Contralateral second dose improves antibody responses to a two-dose mRNA vaccination regimen. J Clin Invest. 10.1172/JCI176411.

24. Jiang, W., Maldeney, A.R., Yuan, X., Richer, M.J., Renshaw, S.E., and Luo, W. (2024). Ipsilateral immunization after a prior SARS-CoV-2 mRNA vaccination elicits superior B cell responses compared to contralateral immunization. Cell Rep 43, 113665. 10.1016/j.celrep.2023.113665.

25. Corbett, K.S., Edwards, D., Leist, S.R., Abiona, O.M., Boyoglu-Barnum, S., Gillespie, R.A., Himansu, S., Schafer, A., Ziwawo, C.T., DiPiazza, A.T., et al. (2020). SARS-CoV-2 mRNA Vaccine Development Enabled by Prototype Pathogen Preparedness. bioRxiv. 10.1101/2020.06.11.145920.

26. Case, J.B., Rothlauf, P.W., Chen, R.E., Liu, Z., Zhao, H., Kim, A.S., Bloyet, L.M., Zeng, Q., Tahan, S., Droit, L., et al. (2020). Neutralizing Antibody and Soluble ACE2 Inhibition of a Replication-Competent VSV-SARS-CoV-2 and a Clinical Isolate of SARS-CoV-2. Cell Host Microbe 28, 475–485 e475. 10.1016/j.chom.2020.06.021.

27. Khoury, D.S., Cromer, D., Reynaldi, A., Schlub, T.E., Wheatley, A.K., Juno, J.A., Subbarao, K., Kent, S.J., Triccas, J.A., and Davenport, M.P. (2021). Neutralizing antibody levels are highly predictive of immune protection from symptomatic SARS-CoV-2 infection. Nat Med 27, 1205–1211. 10.1038/s41591-021-01377-8.

28. Fang, Z., Peng, L., Filler, R., Suzuki, K., McNamara, A., Lin, Q., Renauer, P.A., Yang, L., Menasche, B., Sanchez, A., et al. (2022). Omicron-specific mRNA vaccination alone and as a heterologous booster against SARS-CoV-2. bioRxiv. 10.1101/2022.02.14.480449.

29. Ying, B., Darling, T.L., Desai, P., Liang, C.Y., Dmitriev, I.P., Soudani, N., Bricker, T., Kashentseva, E.A., Harastani, H., Raju, S., et al. (2024). Mucosal vaccine-induced cross-reactive CD8(+) T cells protect against SARS-CoV-2 XBB.1.5 respiratory tract infection. Nat Immunol 25, 537–551. 10.1038/s41590-024-01743-x.

30. Liu, J., Yu, J., McMahan, K., Jacob-Dolan, C., He, X., Giffin, V., Wu, C., Sciacca, M., Powers, O., Nampanya, F., et al. (2022). CD8 T cells contribute to vaccine protection against SARS-CoV-2 in macaques. Sci Immunol 7, eabq7647. 10.1126/sciimmunol.abq7647.

31. Moss, P. (2022). The T cell immune response against SARS-CoV-2. Nat Immunol 23, 186–193. 10.1038/s41590-021-01122-w.

32. McMahan, K., Yu, J., Mercado, N.B., Loos, C., Tostanoski, L.H., Chandrashekar, A., Liu, J., Peter, L., Atyeo, C., Zhu, A., et al. (2021). Correlates of protection against SARS-CoV-2 in rhesus macaques. Nature 590, 630–634. 10.1038/s41586-020-03041-6.

33. Crotty, S. (2019). T Follicular Helper Cell Biology: A Decade of Discovery and Diseases. Immunity 50, 1132–1148. 10.1016/j.immuni.2019.04.011.

34. Halfmann, P.J., Iida, S., Iwatsuki-Horimoto, K., Maemura, T., Kiso, M., Scheaffer, S.M., Darling, T.L., Joshi, A., Loeber, S., Singh, G., et al. (2022). SARS-CoV-2 Omicron virus causes attenuated disease in mice and hamsters. Nature. 10.1038/s41586-022-04441-6.

35. Bentley, E.G., Kirby, A., Sharma, P., Kipar, A., Mega, D.F., Bramwell, C., Penrice-Randal, R., Prince, T., Brown, J.C., Zhou, J., et al. (2021). SARS-CoV-2 Omicron-B.1.1.529 Variant leads to less severe disease than Pango B and Delta variants strains in a mouse model of severe COVID-19. bioRxiv, 2021.2012.2026.474085. 10.1101/2021.12.26.474085.

36. Shuai, H., Chan, J.F., Hu, B., Chai, Y., Yuen, T.T., Yin, F., Huang, X., Yoon, C., Hu, J.C., Liu, H., et al. (2022). Attenuated replication and pathogenicity of SARS-CoV-2 B.1.1.529 Omicron. Nature. 10.1038/s41586-022-04442-5.

37. Yoshihiro, K., Ryuta, U., Maki, K., Shun, I., Masaki, I., Emi, T., Makoto, K., Peter, H., Samantha, L., Tadashi, M., et al. (2022). Characterization and antiviral susceptibility of SARS-CoV-2 Omicron/BA.2. Nature Portfolio. 10.21203/rs.3.rs-1375091/v1.

38. Mesin, L., Schiepers, A., Ersching, J., Barbulescu, A., Cavazzoni, C.B., Angelini, A., Okada, T., Kurosaki, T., and Victora, G.D. (2020). Restricted Clonality and Limited Germinal Center Reentry Characterize Memory B Cell Reactivation by Boosting. Cell 180, 92–106 e111. 10.1016/j.cell.2019.11.032.

39. Viant, C., Weymar, G.H.J., Escolano, A., Chen, S., Hartweger, H., Cipolla, M., Gazumyan, A., and Nussenzweig, M.C. (2020). Antibody Affinity Shapes the Choice between Memory and Germinal Center B Cell Fates. Cell 183, 1298–1311 e1211. 10.1016/j.cell.2020.09.063.

40. Laczko, D., Hogan, M.J., Toulmin, S.A., Hicks, P., Lederer, K., Gaudette, B.T., Castano, D., Amanat, F., Muramatsu, H., Oguin, T.H., 3rd, et al. (2020). A Single Immunization with Nucleoside-Modified mRNA Vaccines Elicits Strong Cellular and Humoral Immune Responses against SARS-CoV-2 in Mice. Immunity 53, 724–732 e727. 10.1016/j.immuni.2020.07.019.

41. Li, C., Lee, A., Grigoryan, L., Arunachalam, P.S., Scott, M.K.D., Trisal, M., Wimmers, F., Sanyal, M., Weidenbacher, P.A., Feng, Y., et al. (2022). Mechanisms of innate and adaptive immunity to the Pfizer-BioNTech BNT162b2 vaccine. Nat Immunol 23, 543–555. 10.1038/s41590-022-01163-9.

42. Lederer, K., Castano, D., Gomez Atria, D., Oguin, T.H., 3rd, Wang, S., Manzoni, T.B., Muramatsu, H., Hogan, M.J., Amanat, F., Cherubin, P., et al. (2020). SARS-CoV-2 mRNA Vaccines Foster Potent Antigen-Specific Germinal Center Responses Associated with Neutralizing Antibody Generation. Immunity 53, 1281–1295 e1285. 10.1016/j.immuni.2020.11.009.

43. Mudd, P.A., Minervina, A.A., Pogorelyy, M.V., Turner, J.S., Kim, W., Kalaidina, E., Petersen, J., Schmitz, A.J., Lei, T., Haile, A., et al. (2022). SARS-CoV-2 mRNA vaccination elicits a robust and persistent T follicular helper cell response in humans. Cell 185, 603–613 e615. 10.1016/j.cell.2021.12.026.

44. Muecksch, F., Wang, Z., Cho, A., Gaebler, C., Ben Tanfous, T., DaSilva, J., Bednarski, E., Ramos, V., Zong, S., Johnson, B., et al. (2022). Increased memory B cell potency and breadth after a SARS-CoV-2 mRNA boost. Nature 607, 128–134. 10.1038/s41586-022-04778-y.

45. Alsoussi, W.B., Malladi, S.K., Zhou, J.Q., Liu, Z., Ying, B., Kim, W., Schmitz, A.J., Lei, T., Horvath, S.C., Sturtz, A.J., et al. (2023). SARS-CoV-2 Omicron boosting induces de novo B cell response in humans. Nature 617, 592–598. 10.1038/s41586-023-06025-4.

46. Payne, R.P., Longet, S., Austin, J.A., Skelly, D.T., Dejnirattisai, W., Adele, S., Meardon, N., Faustini, S., Al-Taei, S., Moore, S.C., et al. (2021). Immunogenicity of standard and extended dosing intervals of BNT162b2 mRNA vaccine. Cell 184, 5699–5714 e5611. 10.1016/j.cell.2021.10.011.

47. Hall, V.G., Ferreira, V.H., Wood, H., Ierullo, M., Majchrzak-Kita, B., Manguiat, K., Robinson, A., Kulasingam, V., Humar, A., and Kumar, D. (2022). Delayed-interval BNT162b2 mRNA COVID-19 vaccination enhances humoral immunity and induces robust T cell responses. Nat Immunol 23, 380–385. 10.1038/s41590-021-01126-6.

48. Baine, Y., and Thorbecke, G.J. (1982). Induction and persistence of local B cell memory in mice. J Immunol 128, 639–643.

49. Kim, J.H., Davis, W.G., Sambhara, S., and Jacob, J. (2012). Strategies to alleviate original antigenic sin responses to influenza viruses. Proc Natl Acad Sci U S A 109, 13751–13756. 10.1073/pnas.0912458109.

50. Kunzli, M., O’Flanagan, S.D., LaRue, M., Talukder, P., Dileepan, T., Stolley, J.M., Soerens, A.G., Quarnstrom, C.F., Wijeyesinghe, S., Ye, Y., et al. (2022). Route of self-amplifying mRNA vaccination modulates the establishment of pulmonary resident memory CD8 and CD4 T cells. Sci Immunol 7, eadd3075. 10.1126/sciimmunol.add3075.

51. Zang, R., Gomez Castro, M.F., McCune, B.T., Zeng, Q., Rothlauf, P.W., Sonnek, N.M., Liu, Z., Brulois, K.F., Wang, X., Greenberg, H.B., et al. (2020). TMPRSS2 and TMPRSS4 promote SARS-CoV-2 infection of human small intestinal enterocytes. Sci Immunol 5. 10.1126/sciimmunol.abc3582.

52. Chen, R.E., Zhang, X., Case, J.B., Winkler, E.S., Liu, Y., VanBlargan, L.A., Liu, J., Errico, J.M., Xie, X., Suryadevara, N., et al. (2021). Resistance of SARS-CoV-2 variants to neutralization by monoclonal and serum-derived polyclonal antibodies. Nat Med 27, 717–726. 10.1038/s41591-021-01294-w.

53. Plante, J.A., Liu, Y., Liu, J., Xia, H., Johnson, B.A., Lokugamage, K.G., Zhang, X., Muruato, A.E., Zou, J., Fontes-Garfias, C.R., et al. (2021). Spike mutation D614G alters SARS-CoV-2 fitness. Nature 592, 116–121. 10.1038/s41586-020-2895-3.

54. Imai, M., Iwatsuki-Horimoto, K., Hatta, M., Loeber, S., Halfmann, P.J., Nakajima, N., Watanabe, T., Ujie, M., Takahashi, K., Ito, M., et al. (2020). Syrian hamsters as a small animal model for SARS-CoV-2 infection and countermeasure development. Proc Natl Acad Sci U S A 117, 16587–16595. 10.1073/pnas.2009799117.

55. Nelson, J., Sorensen, E.W., Mintri, S., Rabideau, A.E., Zheng, W., Besin, G., Khatwani, N., Su, S.V., Miracco, E.J., Issa, W.J., et al. (2020). Impact of mRNA chemistry and manufacturing process on innate immune activation. Science advances 6, eaaz6893. 10.1126/sciadv.aaz6893.

56. Corbett, K.S., Edwards, D.K., Leist, S.R., Abiona, O.M., Boyoglu-Barnum, S., Gillespie, R.A., Himansu, S., Schäfer, A., Ziwawo, C.T., DiPiazza, A.T., et al. (2020). SARS-CoV-2 mRNA vaccine design enabled by prototype pathogen preparedness. Nature 586, 567–571. 10.1038/s41586-020-2622-0.

57. Hassett, K.J., Benenato, K.E., Jacquinet, E., Lee, A., Woods, A., Yuzhakov, O., Himansu, S., Deterling, J., Geilich, B.M., Ketova, T., et al. (2019). Optimization of Lipid Nanoparticles for Intramuscular Administration of mRNA Vaccines. Mol Ther Nucleic Acids 15, 1–11. 10.1016/j.omtn.2019.01.013.

58. Amanat, F., Thapa, M., Lei, T., Ahmed, S.M.S., Adelsberg, D.C., Carreno, J.M., Strohmeier, S., Schmitz, A.J., Zafar, S., Zhou, J.Q., et al. (2021). SARS-CoV-2 mRNA vaccination induces functionally diverse antibodies to NTD, RBD, and S2. Cell 184, 3936–3948 e3910. 10.1016/j.cell.2021.06.005.

59. Stadlbauer, D., Amanat, F., Chromikova, V., Jiang, K., Strohmeier, S., Arunkumar, G.A., Tan, J., Bhavsar, D., Capuano, C., Kirkpatrick, E., et al. (2020). SARS-CoV-2 Seroconversion in Humans: A Detailed Protocol for a Serological Assay, Antigen Production, and Test Setup. Curr Protoc Microbiol 57, e100. 10.1002/cpmc.100.

60. VanBlargan, L., Adams, L., Liu, Z., Chen, R.E., Gilchuk, P., Raju, S., Smith, B., Zhao, H., Case, J.B., Winkler, E.S., et al. (2021). A potently neutralizing anti-SARS-CoV-2 antibody inhibits variants of concern by binding a highly conserved epitope. bioRxiv. 10.1101/2021.04.26.441501.

61. Case, J.B., Bailey, A.L., Kim, A.S., Chen, R.E., and Diamond, M.S. (2020). Growth, detection, quantification, and inactivation of SARS-CoV-2. Virology 548, 39–48. 10.1016/j.virol.2020.05.015.

