## Supplementary figures and images for "Ipsilateral or contralateral boosting of mice with mRNA vaccines confers equivalent immunity and protection against a SARS-CoV-2 Omicron strain"

### Figure S1

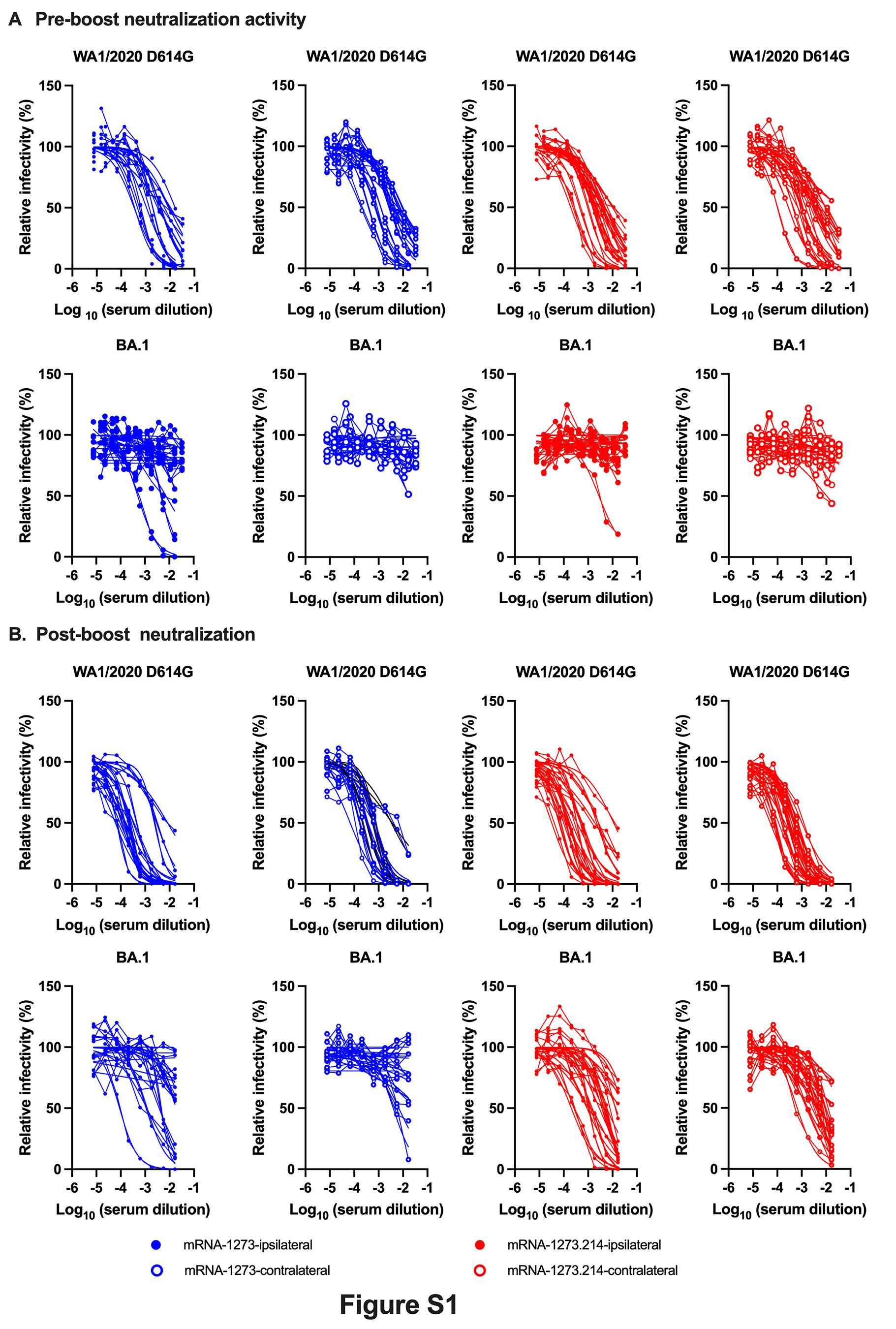

### Figure S2

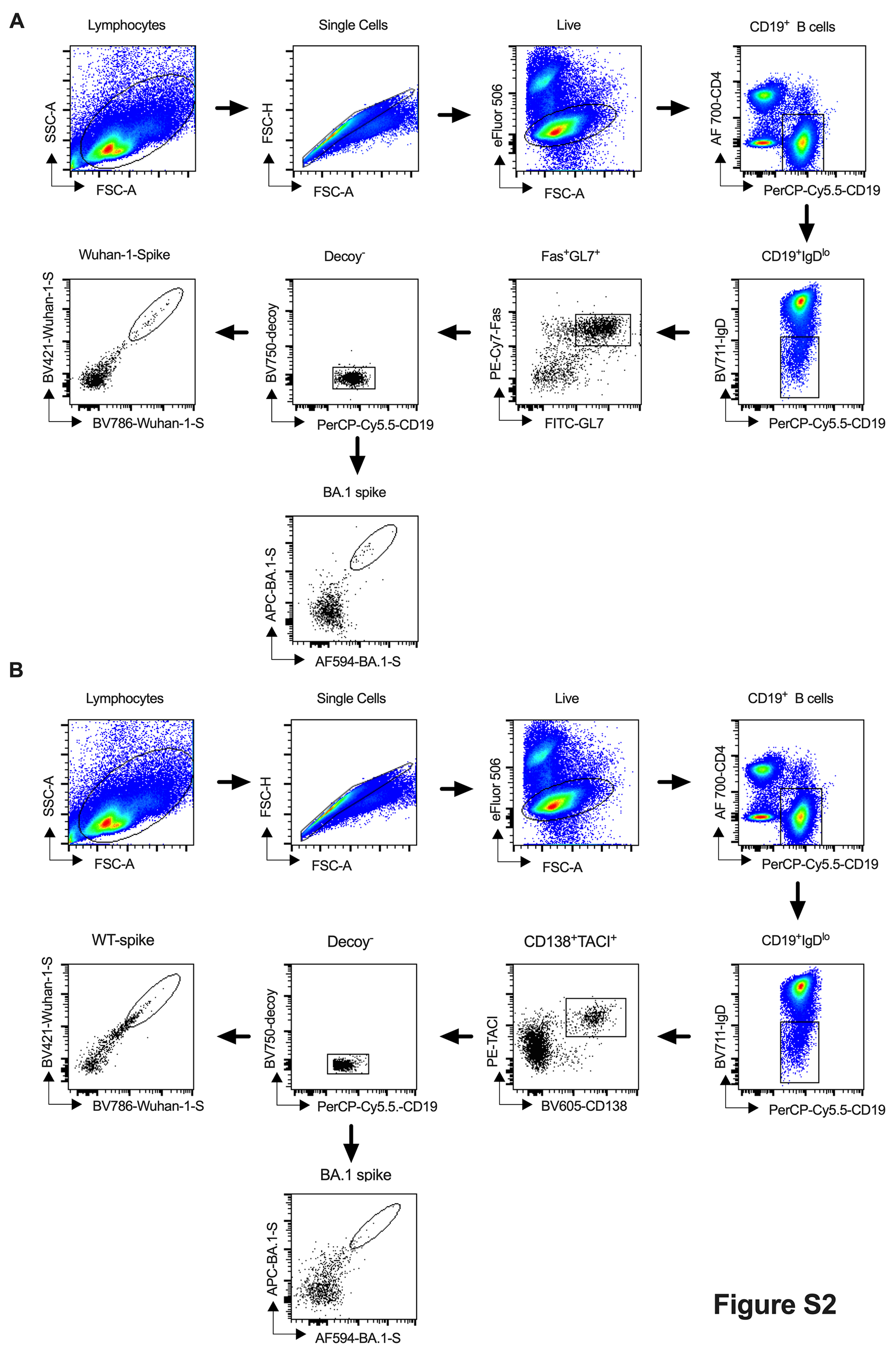

### Figure S3

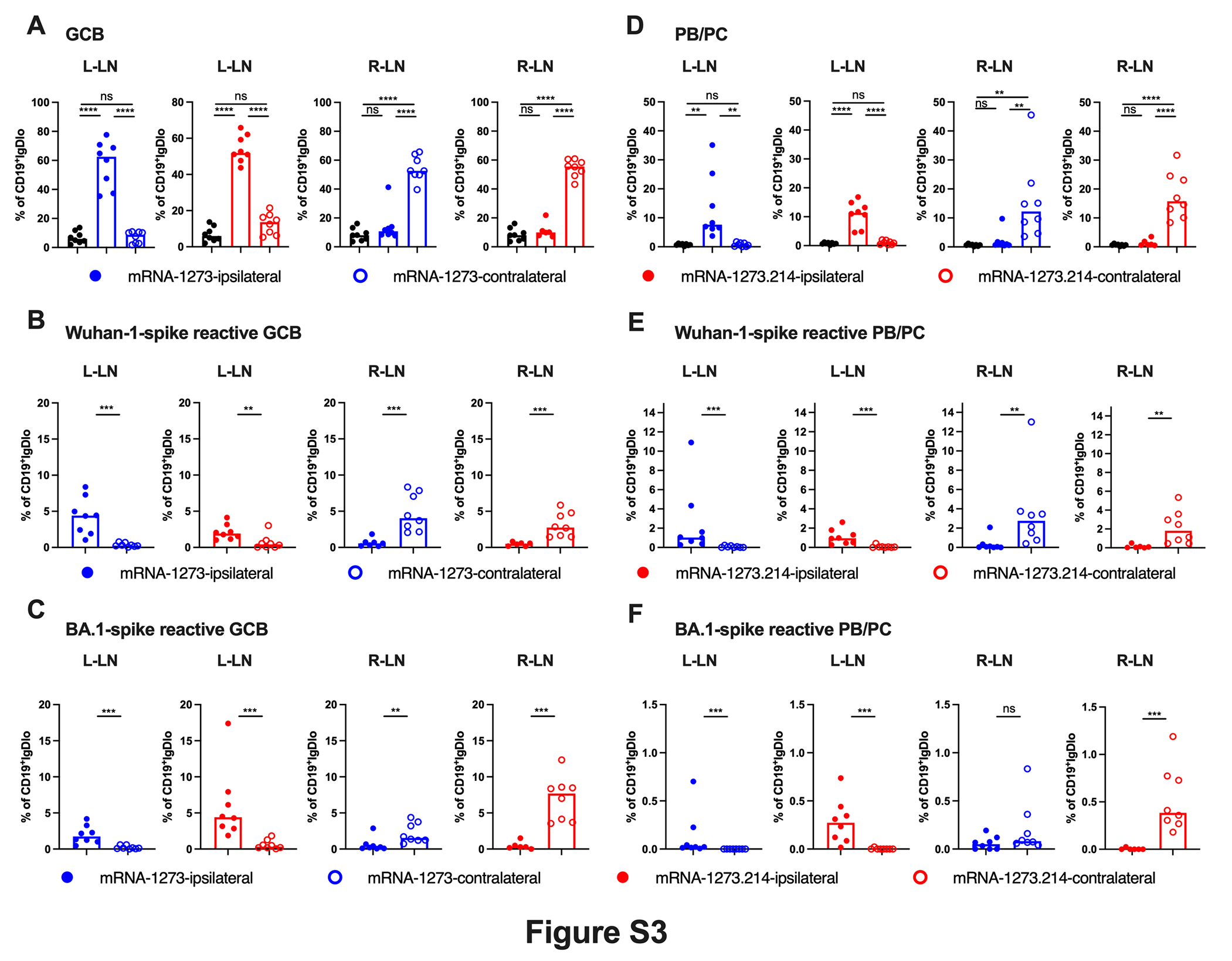

### Figure S4

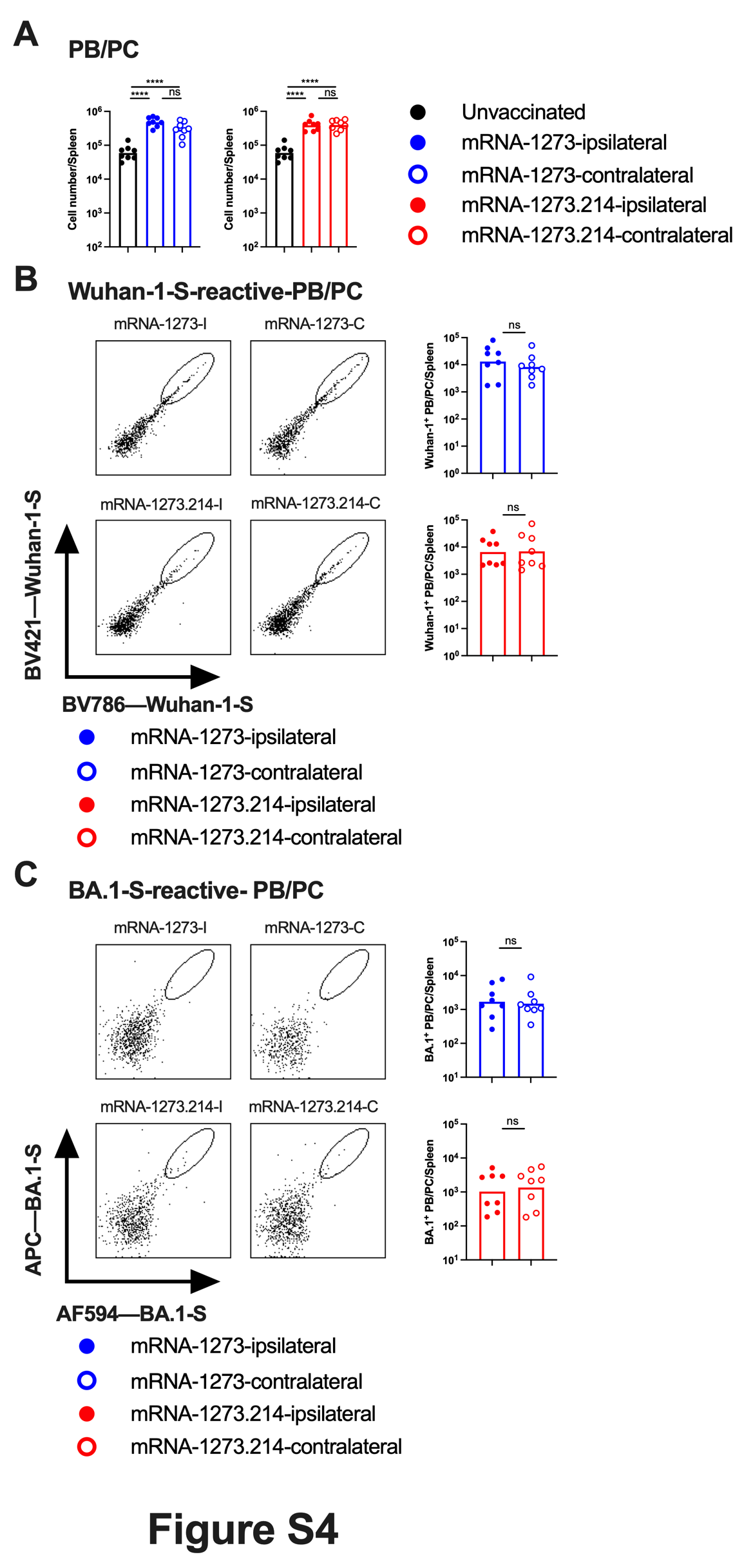

### Figure S5

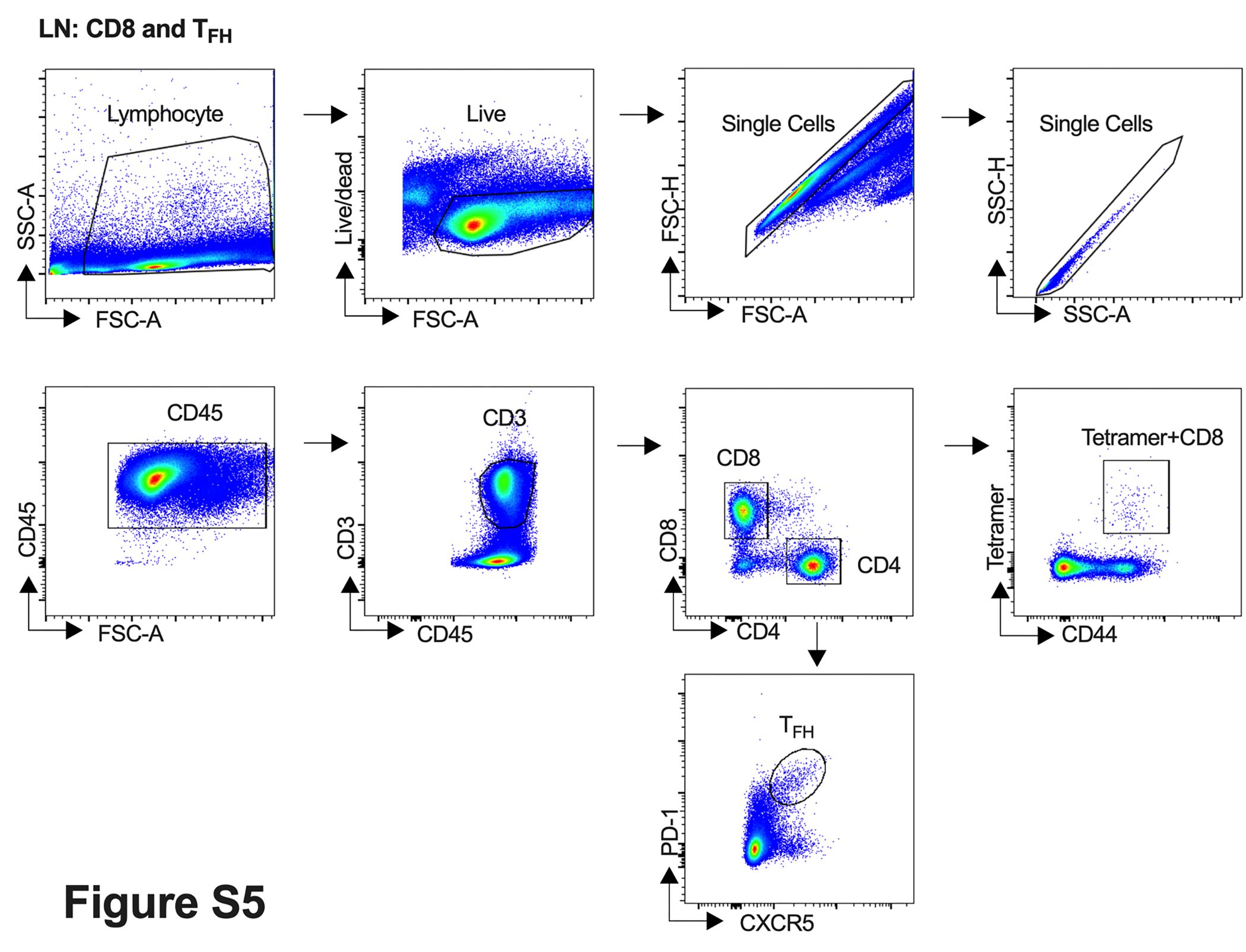

### Figure S6

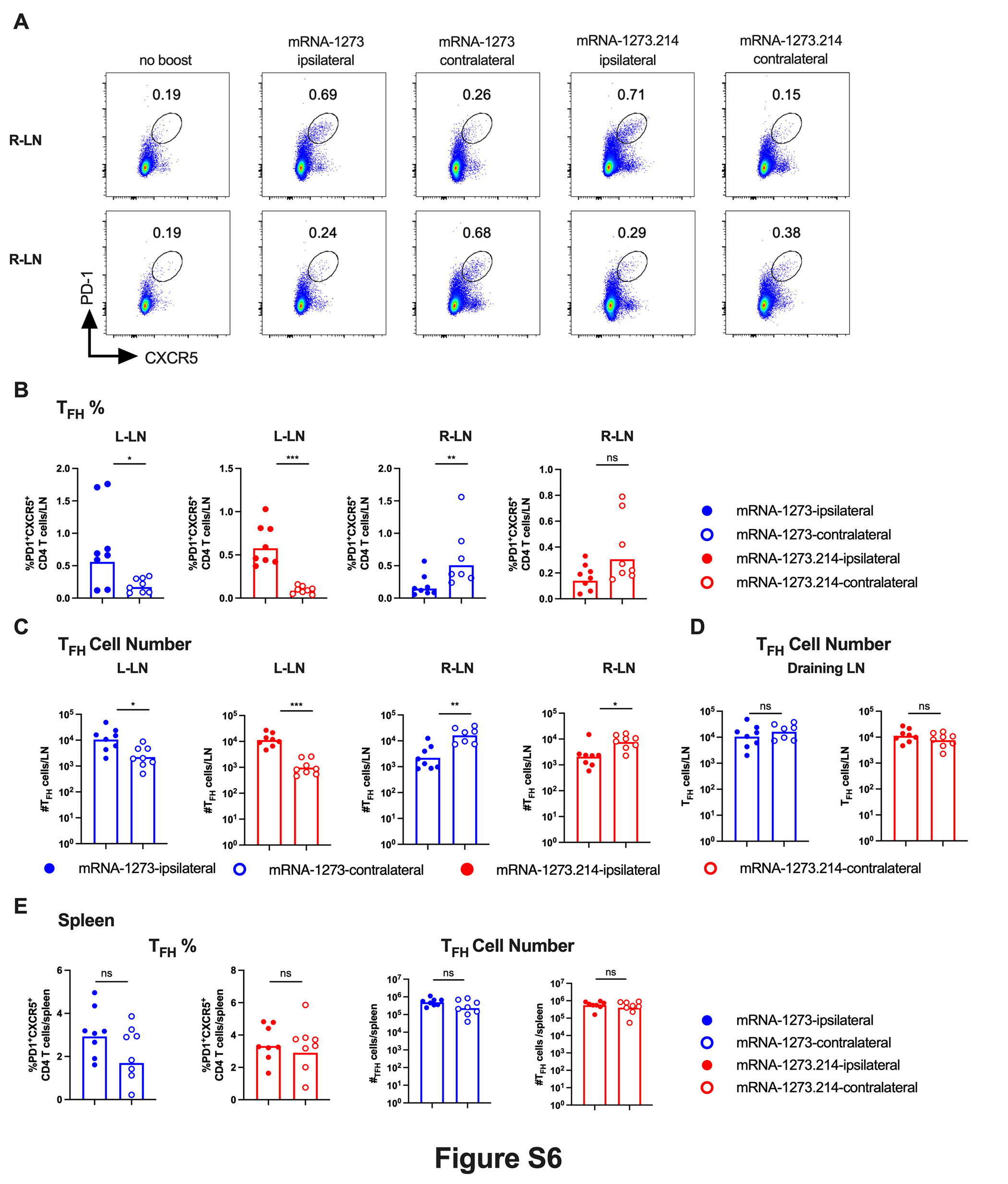
